# The purinergic receptor P2X7 and the NLRP3 inflammasome are druggable host factors required for SARS-CoV-2 infection

**DOI:** 10.1101/2023.04.05.531513

**Authors:** Déborah Lécuyer, Roberta Nardacci, Désirée Tannous, Emie Gutierrez-Mateyron, Aurélia Deva-Nathan, Frédéric Subra, Cristina Di Primio, Paola Quaranta, Vanessa Petit, Clémence Richetta, Ali Mostefa-Kara, Franca Del Nonno, Laura Falasca, Romain Marlin, Pauline Maisonnasse, Julia Delahousse, Juliette Pascaud, Eric Deprez, Marie Naigeon, Nathalie Chaput, Angelo Paci, Véronique Saada, David Ghez, Xavier Mariette, Mario Costa, Mauro Pistello, Awatef Allouch, Olivier Delelis, Mauro Piacentini, Roger Le Grand, Jean-Luc Perfettini

## Abstract

Purinergic receptors and NOD-like receptor protein 3 (NLRP3) inflammasome regulate inflammation and viral infection, but their effects on severe acute respiratory syndrome coronavirus 2 (SARS-CoV-2) infection remain poorly understood. Here, we report that the purinergic receptor P2X7 and NLRP3 inflammasome are cellular host factors required for SARS-CoV-2 infection. Lung autopsies from patients with severe coronavirus disease 2019 (COVID-19) reveal that NLRP3 expression is increased in host cellular targets of SARS-CoV-2 including alveolar macrophages, type II pneumocytes and syncytia arising from the fusion of infected macrophages, thus suggesting a potential role of NLRP3 and associated signaling pathways to both inflammation and viral replication. In vitro studies demonstrate that NLRP3-dependent inflammasome activation is detected upon macrophage abortive infection. More importantly, a weak activation of NLRP3 inflammasome is also detected during the early steps of SARS-CoV-2 infection of epithelial cells and promotes the viral replication in these cells. Interestingly, the purinergic receptor P2X7, which is known to control NLRP3 inflammasome activation, also favors the replication of D614G and alpha SARS-CoV-2 variants. Altogether, our results reveal an unexpected relationship between the purinergic receptor P2X7, the NLRP3 inflammasome and the permissiveness to SARS-CoV-2 infection that offers novel opportunities for COVID-19 treatment.

## Introduction

The rapid worldwide spread of SARS-CoV-2 poses a global health emergency that has not been resolved. Despite the use of preventive COVID-19 vaccination, neutralizing monoclonal antibodies, steroid or IL-6-targeted treatment that drastically reduced the risk of severe symptoms and death due to SAR-CoV-2 infection (1–6), people who are vaccinated or treated, may still get infected with SARS-CoV-2. Thus, there is an urgent need to combine anti-inflammatory therapies with highly effective antivirals for treating COVID-19. Three antiviral compounds, remdesivir (7), molnupiravir (8, 9), and nirmatrelvir (10) were authorized by the US Food and Drug administration for an emergency use for the treatment of COVID-19 (https://www.fda.gov). These drugs are proposed as therapeutic options, alone or in combination with modifiers of inflammatory responses (such as tocilizumab (11), the Janus kinase inhibitor baricitinib (12)) for patients suffering from severe COVID-19 or for those with high risk of severe disease progression. Nevertheless, selection pressures exerted by these antivirals in patients with prolonged infection have been shown to increase viral diversity, to lead to drug resistance and to favor the emergence of mutated variants (13). To prevent further massive outbreaks with emerging variants, a better understanding of mechanisms whereby SARS-CoV-2 hijacks cellular host factors for viral replication and dysregulates anti-viral immune responses is still required and should pave the way for the identification of novel cellular or viral targets, thus offering alternative therapeutic opportunities for SARS-CoV-2-infected patients.

The SARS-CoV-2 entry into host cells begins with the binding of viral spike (S) glycoprotein to angiotensin converting enzyme 2 (ACE-2) and triggers through two distinct pathways, the proteolytic cleavage of S glycoprotein by the serine protease TMPRSS2 at cell surface or by cathepsin B/L in the host cell endosomes (14–16). This process ultimately allows the fusion of viral membranes with host cellular membranes (15, 17) and leads to viral RNA release into host cytosol, to the viral replication using specialized enzymes (such as RNA-dependent RNA polymerase (RdRp) (18)), to viral structural protein expression (such as E and S proteins), and finally, to the assembly and release of the viral progeny (19). Host factors (such as RAB7A, p38MAPK, CK2, AXL and PIFFYVE kinases) are involved in the regulation of early and late steps of SARS-CoV-2 infection (20, 21), but the host cellular pathways used by SARS-COV-2 to establish a viral infection are still poorly understood. Even though SARS-CoV-2-infected people are mainly asymptomatic or exhibit mild to moderate symptoms, approximately 15% of patients experience severe disease with atypical pneumonia and 5% develop an acute respiratory distress syndrome (ARDS) and/or multiple organ failure that is associated with a high mortality rate (around 20-30%) (22). Studies of COVID-19 patients with severe disease revealed a high level of plasma pro-inflammatory cytokines (including IL-β, IL-6, IL-10, IL-18 and TNF) (23) and lactate dehydrogenase (LDH) (24), indicating overt hyper-inflammation during COVID-19, sometimes improperly termed “cytokine storm” or “cytokine release syndrome”. Until now, few molecular mechanisms driving COVID-19 associated hyper-inflammation have been identified and proposed to explain COVID-19 pathogenesis.

SARS-CoV-2 infection activates several microbe-sensing receptors called pattern recognition receptors (PRRs) such as Toll-like receptors (TLRs), RIG-I-like receptors (RLRs) and C-Type lectin receptors (CLRs). Their activations lead to antiviral immune responses driven by the nuclear factor NF-κB, type I interferon secretion and interferon stimulated genes (ISGs) expression (25). Among those PRRs, nucleotide-binding domain, leucine-rich repeat-containing receptor (NLRs) proteins were widely found to be involved in SARS-CoV-2-mediated hyper-inflammation. NLR proteins and purinergic (P2) receptors are the main germline-encoded pattern recognition receptors regulating the secretion of IL-1 family members in response to microbial infection, inflammation, and inflammatory diseases. Upon activation, NLR protein 3 (NLRP3), which is the most studied NLR protein (26), forms large complexes with the adaptor protein ASC (apoptosis-associated speck-like protein containing a CARD domain), called inflammasomes, which activate caspase-1, induce the release of mature cytokines IL-1β and IL-18 (26, 27) and can lead to the inflammatory cell death of stimulated, stressed or infected host cells, which is also known as pyroptosis (28). High levels of IL-1β, IL-18 and LDH positively correlate with disease severity in COVID-19 patients (29), suggesting that inflammasomes participate in SARS-CoV-2 pathogenesis. More recently, NLRP3 inflammasome was found activated in response to SARS-CoV-2 infection and identified as critical driver of COVID-19 (30). In addition, SARS-CoV-2 viral proteins such as viral spike (S) glycoprotein (31), SARS-CoV open reading frame-8b (ORF8b) (32) and the transmembrane pore-forming viral Viroporin 3a (also known as SARS-COV 3a) (33) were shown to activate the NLRP3 inflammasome (30), thus revealing that NLRP3 inflammasome is potentially an interesting molecular target for the treatment of COVID-19. Purinergic receptors are membrane-bound innate receptors that bind extracellular nucleotides (such as adenosine triphosphate (ATP)), and control numerous cellular functions (such as cytokine secretion and migration) mainly in immune cells, but also on other cell types that are involved in SARS-CoV-2 pathogenesis such as type 1 and 2 pneumocytes, endothelial cells, platelets, cardiomyocytes and kidney cells (34, 35). Purinergic P2 receptors are divided into two families, the ionotropic P2X receptors and the metabotropic P2Y receptors, which can regulate the NLRP3 inflammasome (35–37). P2X7 activation was extensively shown to control NLRP3 inflammasome activation and cytokine release in response to danger signals (38). We previously revealed that purinergic receptors (such as P2Y2 and P2X7), NLRP3 inflammasome and associated signaling pathways also control viral replication through the modulation of the fusogenic activity of HIV-1 envelope (37, 39, 40) or the pre-integrative steps of HIV-1 life cycle (41). The contribution of purinergic receptors to viral infection has been confirmed with other purinergic receptors and with several viruses (such as human cytomegalovirus and hepatitis B, C and D viruses) (42). In this context, we hypothesized that the purinergic receptor P2X7 and NLRP3 inflammasome activation could contribute to SARS-CoV-2 pathogenesis and thus, the therapeutic modulation of P2X7 and NLRP3 could regulate the permissiveness of host cells to SARS-CoV-2 infection.

## Results

### Increased NLRP3 expression level in alveolar macrophages, pneumocytes and syncytia detected in lung autopsies of patients with SARS-CoV-2 pneumonia

To study the potential contribution of NLRP3 to SARS-CoV-2 pathogenesis, we first analyzed the expression of NLRP3 in lung tissue samples obtained from three uninfected carriers and seven SARS-CoV-2-infected patients who died from COVID-19. Autopsies were stained with hematoxylin and eosin, or incubated with antibody against NLRP3 and analyzed. As previously described (43), parenchymal multifocal damages with intra-alveolar inflammation, fibrin and hyaline membrane formation that are consistent with a diagnosis of diffuse alveolar damage were observed in all SARS-CoV-2-infected carriers. The pattern of developed pneumonia (with fibrotic organization and type 2 pneumocyte hyperplasia) and fibroblastic foci formed by the loss of organized connective tissue consistent with alveolar duct fibrosis were also detected (Fig. 1A). Besides, the presence of inflammatory cells (composed mainly of macrophages and lymphocytes) was the main characteristic of COVID-19 patient autopsies (Fig. 1A). NLRP3 expression was then examined on these samples, and immuno-reactive NLRP3 was mainly detected in Type II pneumocytes (Fig. 1, B and C) and alveolar macrophages (Fig. 1, B and D) from both SARS-CoV-2 patients and uninfected specimens, which have been previously shown to be cellular targets of SARS-CoV-2 (44, 45). Interestingly, we also detected in SARS-CoV-2-infected lung tissues, some syncytia that expressed macrophage marker, CD68 (Fig. 1E), NLRP3 (Fig. 1F) and viral RNA (Fig. 1G), revealing a potential link between NLRP3 expression and viral infection.

**Fig. 1.**
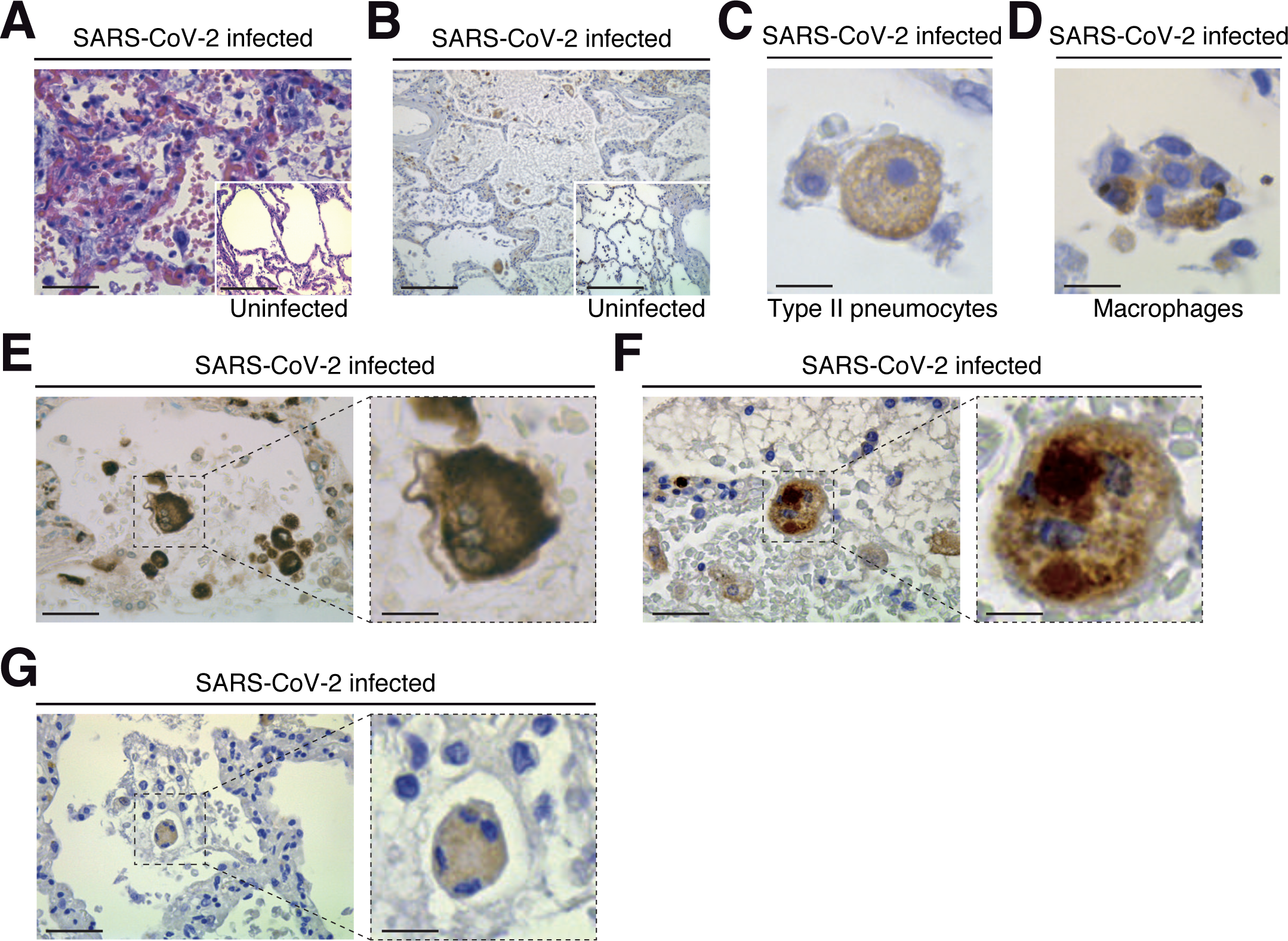
Pathological changes and NLRP3 expression in the lungs of COVID-19 patients with severe disease. **A** Parenchymal damage (with inflammation and fibrous proliferative phase) and hyperplasic amphophilic type II pneumocytes are detected in COVID-19 patients as compared to non-COVID-19 patients (insert). Hematoxylin and eosin staining is shown (bars, 22 µm and 50 µm). **B** NLRP3 expression is detected into alveolar septa and lumen in both COVID-19 and non-COVID-19 patients (bars, 45 µm and 50 µm). **C** Type II pneumocytes with cytoplasmic NLRP3 expression (bar, 3 µm). **D** NLRP3 positive alveolar macrophage in lumen (bar, 3 µm). **E-G** Syncytium detected on COVID-19 patients express macrophage marker CD68 (**E**) (bars, 8 µm and 3 µm), NLRP3 (**F**) (bars, 8 µm and 3 µm) and cytoplasmic double strand RNA, indicative of viral infection (**G**) (bars, 12 µm and 4 µm). Positive stainings (brown) for NLRP3, CD68 or double strand RNA are shown in **B-G**. Magnifications are shown in **E-G**.

### Increased IL-1β secretion is associated with NLRP3 inflammasome activation during abortive macrophage infection with SARS-CoV-2

Given that macrophages are key cellular players for COVID-19 pathogenesis (46) and inflammasome activation was detected in SARS-CoV-2-infected macrophages (30), we first analyzed the expression of NLRP3 during the infection of macrophages with SARS-CoV-2. Interestingly, we found that NLRP3 expression level increased 24 hours after SARS-CoV-2 infection in phorbol-12-myristate-13-acetate (PMA) – differentiated human THP1 macrophages with different multiplicities of infection (MOI) (Fig. 2A and Supplementary Fig. 1A) or at different time points after infection with a MOI of 0.2 (Fig. 2B and Supplementary Fig. 1B). Increased NLRP3 expression level was observed in the absence of detectable intracellular SARS-CoV-2 spike (S) protein expression, indicating a lack of viral replication (Fig. 2, A and B). These results thus confirm, as previously published (30), that NLRP3 expression level is increased during the abortive SARS-CoV-2 infection in macrophages. Then, we evaluated the effect of SARS-CoV-2 infection on two hallmarks of inflammasome activation, the oligomerization of the adaptor protein ASC and the production of mature IL-1β (47). The oligomerization of ASC, revealed by the quantification of ASC speck formation (Fig. 2, C and D), and the release of mature IL-1β (Fig. 2, E and F) were increased in PMA-THP1 macrophages that were infected with SARS-CoV-2, as compared to uninfected cells (Fig. 2, C-F). Inactivation of NLRP3 inflammasome with a pharmacological inhibitor MCC950 (Fig. 2G and Supplementary Fig. 1C) or using specific NLRP3 CRISPR guide RNAs (gRNAs) and the *CAS9* gene (CrNLRP3) (Fig. 2H and Supplementary Fig. 1D) significantly reduced the release of mature IL-1β by PMA-THP1 macrophages that were infected with SARS-CoV-2 during 24 hours (Fig. 2, I and J). These results reveal that NLRP3 inflammasome is activated during abortive SARS-CoV-2 infection in macrophages. Given that the purinergic receptor P2X7 has been extensively involved in the activation of NLRP3 inflammasome (47), we also investigated the impact of P2X7 antagonist oxidized ATP (OxATP) or agonist 2’(3’)-O-(4-Benzoylbenzoyl) adenosine 5’-triphosphate (BzATP) on NLRP3 inflammasome activation elicited by SARS-CoV-2. Pharmacological inhibition or activation of P2X7 with respectively, OxATP or BzATP (Fig. 2, K and L), did not modulate increased expression level of NLRP3 (Fig. 2K and Supplementary Fig. 1E) nor IL-1β release (Fig. 2L) from PMA-treated THP1 macrophages that were infected with SARS-CoV-2. Results reveal that the activation of NLRP3 inflammasome detected during the infection of PMA-treated THP1 macrophages with SARS-CoV-2 does not require the purinergic receptor P2X7. Taken together, these results indicate that NLRP3 inflammasome activation is detected during the early steps of human macrophage infection with SARS-CoV-2.

**Fig. 2.**
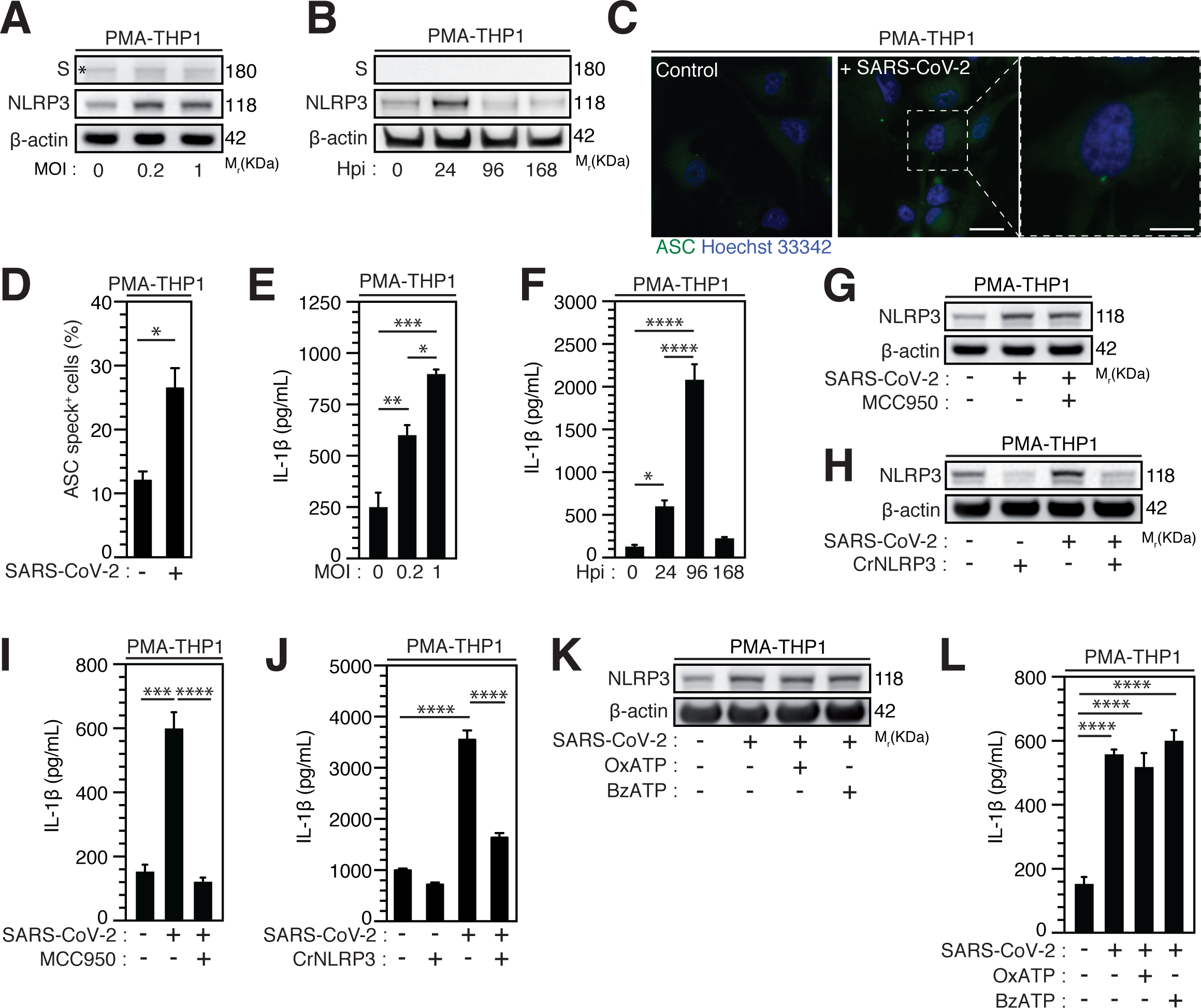
SARS-CoV-2 infection induces NLRP3 inflammasome activation independently of P2X7 in macrophages. **A-F** PMA-differentiated THP1 cells were infected with SARS-CoV-2 for 24 hours at indicated multiplicity of infection (MOI) (**A**) or during indicated hours post-infection (Hpi) (MOI = 0.2) (**B**) and evaluated for Spike (S), NLRP3 and β-actin expression (**A, B**) or IL-1β secretion (**E, F**). Representative images (bars, 20 µm and 10 µm, from left to right) (**C)** and percentages (**D**) of control or PMA-differentiated THP1 cells infected for 24 hours (MOI = 0.2), with ASC speck formation (ASC^+^) are shown. DNA is detected using Hoechst 33342. (**C**, **D**). **G-J** PMA-differentiated THP1 cells that were incubated with the NLRP3 inflammasome inhibitor, MCC950 (20 µM) (**G, I**), and PMA-differentiated, control (CrCo.) or NLRP3-depleted (CrNLRP3) THP1 cells (**H, J**) were infected with SARS-CoV-2 for 24 hours (MOI = 0.2) and evaluated for NLRP3 and β-actin expression (**G, H**) or IL-1β secretion (**I, J**). **K, L** PMA-differentiated THP1 cells were infected with SARS-CoV-2 (MOI = 0.2) for 24 hours, in presence of control vehicle, the P2X7 inhibitor OxATP, or the P2X7 activator BzATP, and evaluated for NLRP3 and β-actin expression (**K**) or IL-1β secretion (**L**). In (**A**), the asterisk (*) is indicating a non-specific band. Data are presented as means ± SEM from at least 3 independent experiments. *p* values (\**p* < 0.05, \*\**p* < 0.01, \*\*\**p* < 0.001 and \*\*\*\**p* < 0.0001) were determined using unpaired t-test (**D**) and one-way ANOVA Tukey’s multiple comparisons test (**E, F, I, J** and **L**).

### NLRP3 acts as a proviral host factor for SARS-CoV-2 infection

Considering that epithelial cells, which have been identified as primary targets of viral replication in the lung and the nasal epithelium of SARS-CoV-2-infected patients (44, 48, 49), also express NLRP3 inflammasome (50), we then analyzed the activation of NLRP3 inflammasome during the infection of permissive host Caco-2 cells and ACE2-overexpressing A549 (ACE2-A549) cells. Surprisingly, a weak, but significant SARS-CoV-2-MOI-dependent release of IL-1β was detected in response to Caco-2 (Fig. 3, A) and ACE2-A549 (Fig. 3, C) infection during 24 hours. The IL-1β secretion was also detected at a MOI of 4 up to 48 hours after infection with SARS-CoV-2 at the same target cells (Fig. 3, B and D). IL-1β secretion is reduced when SARS-CoV-2 was preincubated and neutralized with COVID-19 patient’s serum before infection of ACE2-A549 cells (Fig. 3E), thus revealing that NLRP3 inflammasome is activated after viral entry into permissive host cells. We next evaluated whether the activation of NLRP3 inflammasome in epithelial cells might affect viral replication, as previously shown during SARS-CoV-2 infection of macrophages (30). We determined the impact of MCC950 and another NLRP3 inhibitor, Tranilast, on viral replication. Despite the fact MCC950 did not modulate viral replication in ACE2-A549 cells (Fig. 3K and Supplementary Fig. 2, A-C), Tranilast significantly reduced the replication of SARS-CoV-2 in Caco-2 cells (Fig. 3, F and H and Supplementary Fig. 2, D) and ACE2-A549 cells (Fig. 3, G, I-K and Supplementary Fig. 2, E). These results were revealed by detecting the intracellular expression of spike (S) protein (Fig. 3, F and G and Supplementary Fig. 2, A, B, D and E), the detection of viral E RNA expression (Fig. 3, H and I), the frequency of spike (S)-positive cells (Fig. 3, J and K and Supplementary Fig. 2, C) and the cellular viability of treated cells (Supplementary Fig. 2, F-H). As expected, SARS-CoV-2 that was preincubated with serum of convalescent COVID-19 patients did not infect host cells (Fig. 3, J and K). We also examined the impact of NLRP3 depletion on viral replication using small interfering RNA (Fig. 3, L-O). Importantly, the depletion of NLRP3 in Caco-2 and ACE2-A549 cells (Fig. 3, L and N) decreased the replication of SARS-CoV-2, as revealed by the decrease of viral RdRp RNA (Fig. 3, M and O). Altogether, these results demonstrate that NLRP3 acts as a proviral host factor that is required for SARS-CoV-2 infection and replication in target epithelial cells.

**Fig. 3.**
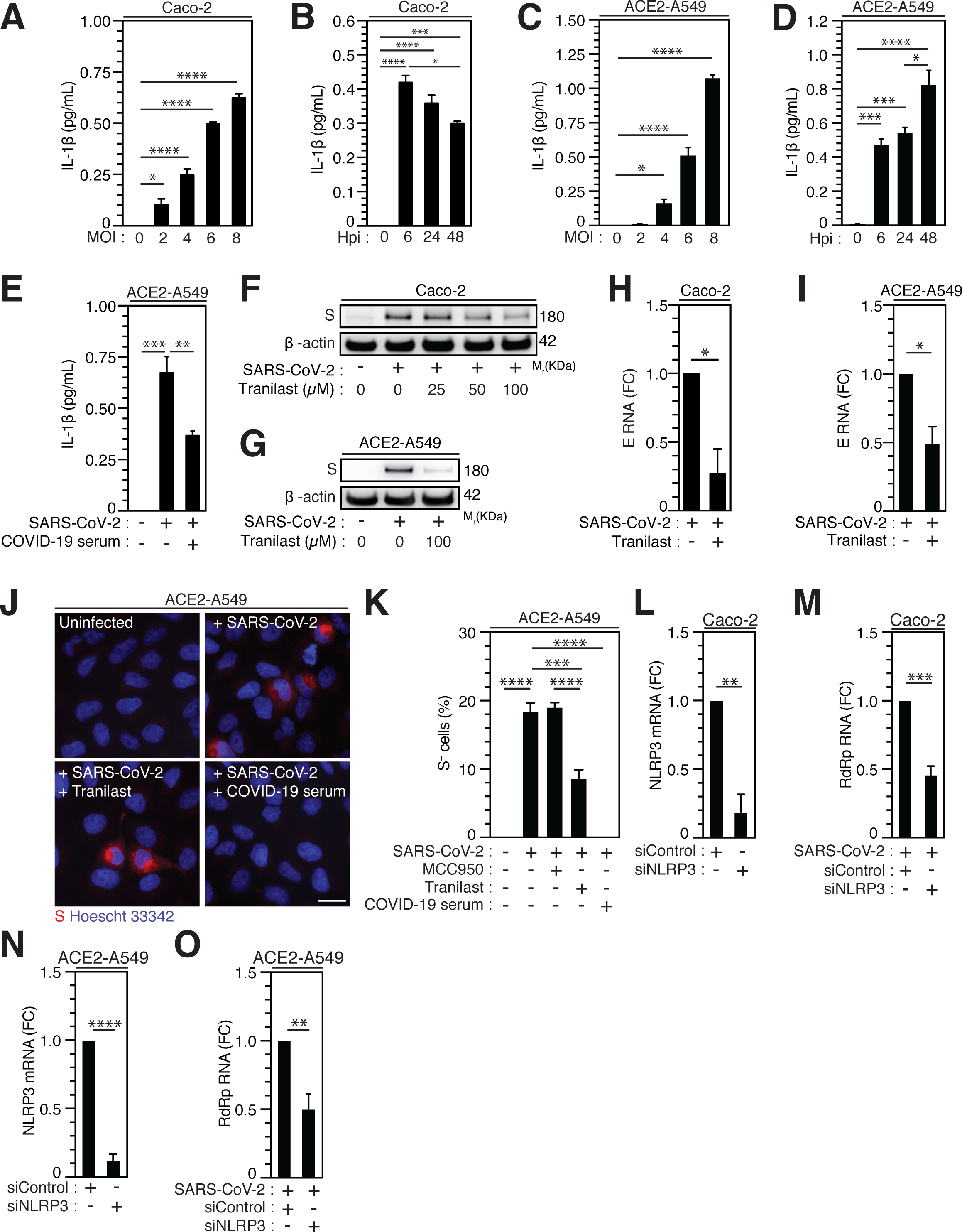
NLRP3 inflammasome activation dictates permissiveness of epithelial cells to SARS-CoV-2 infection. **A-D** Caco-2 (**A, B**) or ACE2-A549 (**C, D**) cells were infected with SARS-CoV-2 for 48 hours at indicated MOI (**A, C**) or during indicated hours post-infection (hpi) (MOI = 4) (**B, D**) and IL-1β secretion was monitored. **E** ACE2-A549 cells were infected during 48 hours with SARS-CoV-2 or with SARS-CoV-2 neutralized with convalescent COVID-19 patient serum. IL-1β secretion was then monitored. **F, G** Caco-2 (**F**) and ACE2-A549 (**G**) cells were infected (or not) with SARS-CoV-2 (MOI = 2) during 48 hours in presence of indicated concentrations of Tranilast and evaluated for Spike (S) and β-actin expressions. **H, I** Caco-2 (**H**) or ACE2-A549 (**I**) cells were infected for 48 hours with SARS-CoV-2 (MOI = 0.2) in presence of 100 µM Tranilast, and analyzed for SARS-CoV-2 E RNA expression using quantitative RT-PCR. Fold changes (FC) are indicated. **J, K** ACE2-A549 cells were infected for 48 hours with SARS-CoV-2 (MOI = 2) in presence of 20 µM MCC950, 100 µM Tranilast or with SARS-CoV-2 neutralized with convalescent COVID-19 patient serum and analyzed for spike (S) expression by fluorescence microscopy. Representative images (bar, 20 µm) (**J**) and percentages of spike (S)-positive (S^+^) cells (**K**) are shown. DNA is detected using Hoechst 33342. **L-O** Caco-2 (**L, M**) or ACE2-A549 (**N, O**) cells were subjected to siRNA against NLRP3 for 48 hours and analyzed for NLRP3 mRNA expression (**L, N**) or infected with SARS-CoV-2 for 48 hours and evaluated for SARS-CoV-2 RdRp RNA expression, by quantitative RT-PCR (**M, O**). RT-PCR samples were first normalized by β-actin mRNA level in each sample, and then normalized to the control condition. The data are presented as means ± SEM from at least 3 independent experiments. *p* values (\**p* < 0.05, \*\**p* < 0.01, \*\*\**p* < 0.001 and \*\*\*\**p* < 0.0001) were determined using one-way ANOVA Tukey’s multiple comparisons test (**A-E** and **K**) and unpaired t-test (**H, I** and **L-O**).

### Caspase-1 activation is also required for SARS-CoV-2 replication

Considering that caspase-1, which is a substrate of NLRP3 inflammasome activation (51, 52) mainly involved in the protection of host cells against microbial infection (53), was also shown to promote host cell survival in response to microbial virulence factors such as *Aeromonas* aerolysin (54), we examined the contribution of caspase-1 to SARS-CoV-2 viral replication. The pharmacological inhibition of caspase-1 using YVAD inhibitor in Caco-2 (Fig. 4, A and C) and ACE2-A549 (Fig. 4, B and D) cells reduced SARS-CoV-2 infection, as revealed by the frequency of spike (S)-positive cells at 24 hours after infection (Fig. 4, A-D), without affecting cellular viability (Supplementary Fig. 3, A and B). Accordingly, the depletion of caspase-1 using small interfering RNA in Caco-2 (Fig. 4, E) and ACE2-A549 (Fig. 4, G) cells also significantly reduced the detection of viral RdRp mRNA (Fig. 4, F) and intracellular spike (S) protein (Fig. 4 H and Supplementary Fig. 3C) in CASP-1-depleted cells, as compared to control cells. Altogether, these results reveal that caspase-1 also contributes to SARS-CoV-2 infection and replication.

**Fig. 4.**
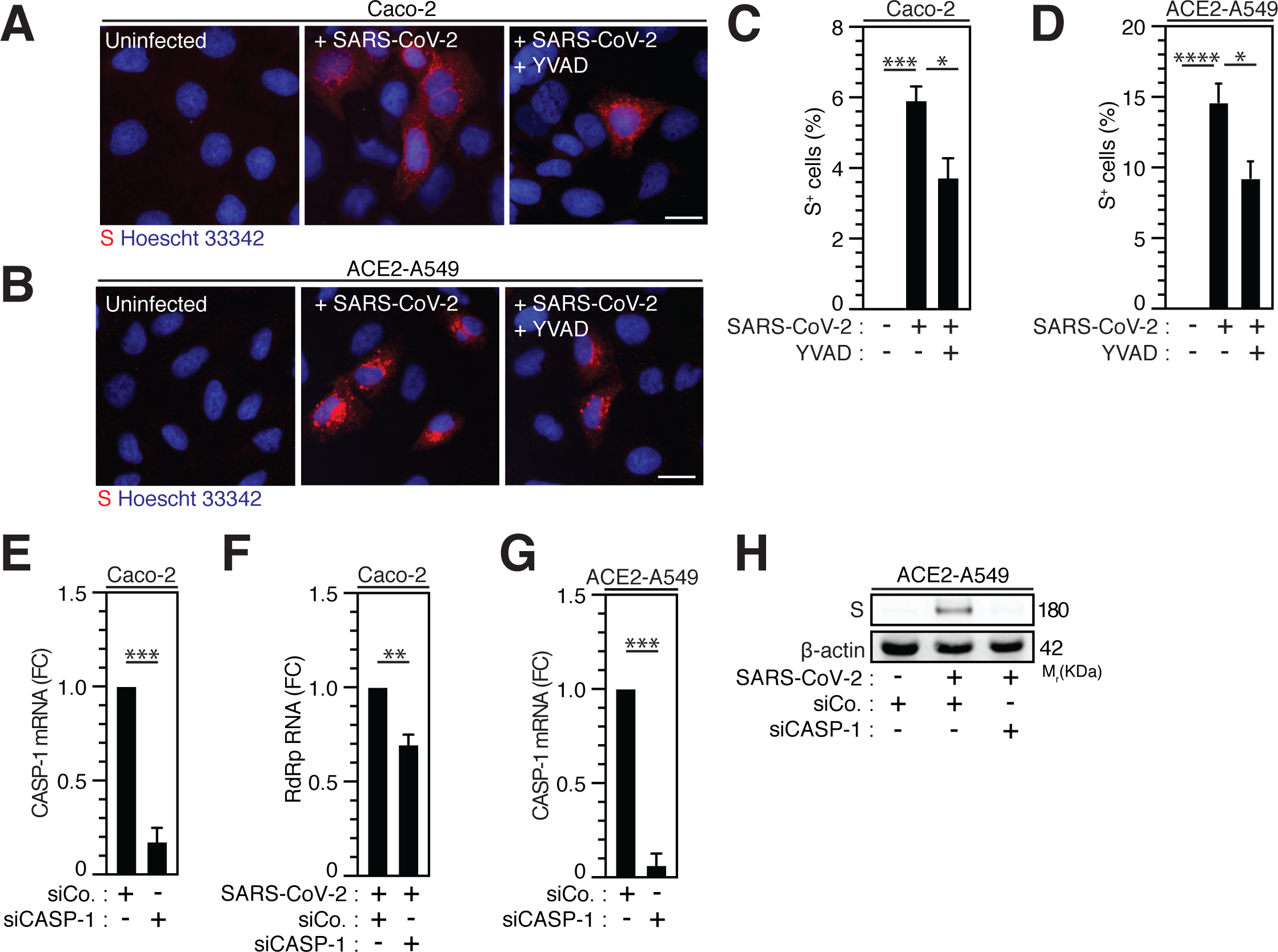
Caspase-1 activity regulates SARS-CoV-2 replication. **A-D** Caco-2 (**A, C**) or ACE2-A549 (**B, D**) were infected with SARS-CoV-2 (MOI = 2) in presence 50 µM of caspase-1 inhibitor, Ac-YVAD-cmk (YVAD), for 48 hours and analyzed for spike (S) expression by fluorescence microscopy. Representative images (bar, 20 µm) (**A, B**) and percentages of spike (S)-positive (S^+^) cells (**C, D**) are shown. DNA is detected using Hoechst 33342. **E-H** Caco-2 (**E, F**) and ACE2-A549 (**G, H**) were treated with siRNA against caspase-1 (CASP-1) for 48 hours and analyzed for CASP-1 mRNA quantification by RT-PCR (**E, G**), or infected with SARS-CoV-2 at MOI = 0.2 (**F**) or MOI = 2 (**H**) during 48 hours, and analyzed for SARS-CoV-2 RdRp RNA expression by quantitative RT-PCR (**F**) or for spike (S) and β-actin expressions (**H**). The data are presented as means ± SEM from at least 3 independent experiments. *p* values (\**p* < 0.05, \*\**p* < 0.01, \*\*\**p* < 0.001 and \*\*\*\**p* < 0.0001) were determined using one-way ANOVA Tukey’s multiple comparisons test (**C, D**) and unpaired t-test (**E-G**).

### Purinergic receptor P2X7 dictates NLRP3 inflammasome activation for SARS-CoV-2 replication

As previously published (37, 39, 55), we initially revealed the contribution of purinergic receptors to the early steps of human immunodeficiency virus-1 (HIV-1) infection. To evaluate the possibility that P2X7, an important partner of NLRP3 inflammasome activation, might also affect SARS-CoV-2 viral replication, we next examined the impact of a non-selective antagonist of purinergic receptors P2X, the pyridoxal-phosphate-6-azopheny-2’,4’-disulfonate (PPADS) and the P2X7-specific antagonist OxATP, on SARS-CoV-2 viral replication and related cellular damage. The inhibition of P2X and P2X7 activities by PPADS and/or OxATP reduced the frequency of spike (S)-positive cells (Fig. 5, A-D) and the expression level of viral E RNA (Fig. 5, E) elicited by SARS-CoV-2 in Vero E6 cells (Fig. 5, A and C) and ACE2-A549 (Fig. 5, B, D and E). PPADS and OxATP partially affected viability of uninfected Vero E6 cells (Supplementary Fig. 4, A and B) at the concentration of 100 µM, but not at working concentration (10 µM). Moreover, OxATP did not show any effect on the viability of ACE2-A549 cells (Supplementary Fig. 4C). Accordingly, P2X7 depletion using small interfering RNA significantly reduced intracellular spike (S) expression level detected after 48-hour infection of ACE2-A549 cells with SARS-CoV-2 (Fig. 5F and Supplementary Fig. 4, D and E). Conversely, the activation of P2X7 with BzATP increased replication of SARS-CoV-2, as revealed by the increase of intracellular spike (S) expression levels (Fig. 5, B and D) and viral E expression level (Fig. 5G), but did not show a significant impact on cellular viability (Supplementary Fig. 4F). We then analyzed impact of OxATP on the replication of SARS-CoV-2 into primary salivary gland epithelial cells, which have been identified as primary targets for SARS-CoV-2 in SARS-CoV-2-infected patients (56), and confirmed that OxATP significantly reduced SARS-CoV-2 replication, as revealed by the reduction of RdRp mRNA expression level in primary salivary gland epithelial cells that were treated with OxATP and infected during 48 hours with SARS-CoV-2 (Fig. 5H). This process occurs without affecting the viability of primary salivary gland epithelial cells (Supplementary Fig. 4G). Finally, we analyzed the effect of P2X7 inhibition on the replication of SARS-CoV-2 Alpha variant and determined the amount of viral RdRp RNA released in the supernatant of liver Huh7 cells that were infected with SARS-CoV-2 Alpha variant in presence or in absence of OxATP. We observed that OxATP significantly reduced the release of viral particles in the supernatant of SARS-CoV-2-infected Huh7 cells (Fig. 5I), thus revealing that purinergic receptor P2X7 also regulates the replication of SARS-CoV-2 Alpha variant. To determine whether the purinergic receptor P2X7 may act upstream of NLRP3 inflammation activation, ACE2-A549 cells were infected during 48 hours with SARS-CoV-2 in presence of indicated concentrations of BzATP and/or Tranilast. As previously shown (Fig. 3, F-K and Fig. 5, B, D and G), the intracellular expression of spike (S) protein elicited by the infection of ACE2-A549 cells with SARS-CoV-2 was increased or decreased in presence of BzATP or Tranilast, respectively (Fig. 5J). Interestingly, we observed that the treatment of ACE2-A549 cells with both BzATP and Tranilast significantly reduced the intracellular spike (S) expression level detected after 48 hours of infection, as compared to ACE2-A549 cells that were infected in presence of BzATP (Fig. 5J), thus demonstrating that the purinergic receptor P2X7 acts upstream to NLRP3 inflammasome activation and dictates viral replication.

**Fig. 5.**
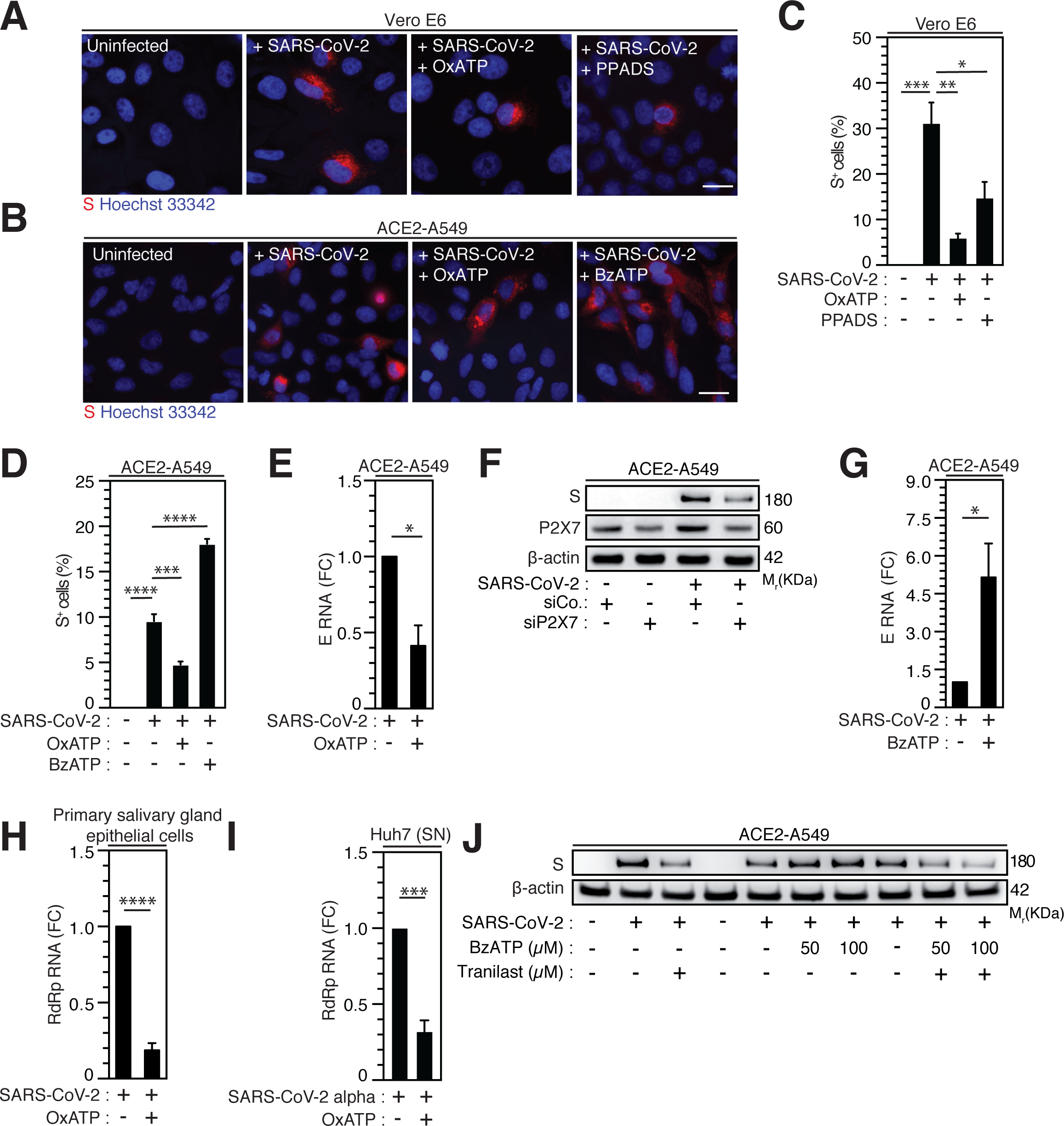
Purinergic receptor P2X7 modulates SARS-CoV-2 replication through NLRP3 inflammasome activation. **A-D** Vero E6 (**A, C**) or ACE2-A549 (**B, D**) cells were infected with SARS-CoV-2 at MOI = 0.1 for 24 hours (**A, C**) or at MOI = 2 for 48 hours (**B, D**), with 10 µM OxATP and 50 µM PPADS (**A, C**) or with 100 µM OxATP and 100 µM BzATP (**B, D**) and analyzed for spike (S) expression by fluorescence microscopy. Representative images (bar, 20 µm) (**A, B**) and percentages of spike (S)-positive (S^+^) cells (**C, D**) are shown. DNA is detected using Hoechst 33342. **E** ACE2-A549 cells were infected with SARS-CoV-2 (MOI = 0.2) for 48 hours in presence of 100 µM OxATP and analyzed for SARS-CoV-2 E RNA expression by quantitative RT-PCR. **F** ACE2-A549 cells were treated with siRNA against P2X7 for 48 hours and analyzed for spike (S), P2X7 and β-actin expression. **G** ACE2-A549 cells were infected with SARS-CoV-2 (MOI = 0.2) for 48 hours in presence of 100 µM BzATP and analyzed for SARS-CoV-2 E RNA expression by quantitative RT-PCR. **H** Primary salivary gland epithelial cells were infected with SARS-CoV-2 (MOI = 0.5) for 48 hours in presence of 100 µM OxATP and analyzed for SARS-CoV-2 RdRp RNA expression by quantitative RT-PCR. **I** Huh7 cells were infected with SARS-CoV-2 Alpha variant in presence of 100 µM OxATP for 48 hours. Then, the viral yielding was determined by detecting RdRp RNA expression by quantitative RT-PCR in the supernatant (SN) of control and SARS-CoV-2 Alpha variant infected Huh7 cells. Fold changes (FC) are indicated in (**E**) and (**G-I**). **J** ACE2-A549 cells were infected with SARS-CoV-2 during 48 hours (MOI = 2) in presence of control, 50 or 100 µM BzATP, 100 µM Tranilast, or the combination of either 50 µM BzATP and 100 µM Tranilast or 100 µM BzATP and 100 µM Tranilast, and analyzed for spike (S) and β-actin expression. All experiments are representative of at least 3 independent experiments. The data are presented as means ± SEM from at least 3 independent experiments. *p* values (\**p* < 0.05, \*\**p* < 0.01, \*\*\**p* < 0.001 and \*\*\*\**p* < 0.0001) were determined using one-way ANOVA Tukey’s multiple comparisons test (**C, D**) and unpaired t-test (**E, G-I**).

### Purinergic receptor P2X7 and NLRP3 inflammasome regulate SARS-CoV-2 replication at a post-entry level

Then, we determined whether purinergic receptor P2X7 and NLRP3 might control SARS-CoV-2 replication through the modulation of viral entry. We analyzed viral entry by using SARS-CoV-2 spike (S) pseudotyped HIV-1. Thus, we incubated ACE2 and transmembrane serine protease 2 (TMPRSS2)-expressing A549 (ACE2-TMPRSS2-A549) highly permissive cells with OxATP, BzATP or serum of convalescent COVID-19 patients during 4 hours before their infection with green fluorescent protein (GFP)-tagged HIV-1NL4-3**_Δ_**_Env_ variant (defective in viral envelope) pseudotyped with the SARS-CoV-2 spike (S) envelope (S-GFP-LV) (Fig. 6, A-E). Then, we analyzed the viral entry by detecting after 48-hour infection, the frequency of GFP^+^ cells and observed that OxATP and BzATP did not reduce the frequency of GFP^+^ cells detected in S-GFP-LV-infected ACE2-TMPRSS2-A549 cells (Fig. 6, A-D and F). Of note, serum of convalescent COVID-19 patients efficiently reduced the frequency of GFP^+^ cells detected after 48 hours of infection (Fig. 6, E and F). These results were confirmed by detecting the intracellular HIV-1 Cap24 capsid in ACE2-TMPRSS2-A549 cells (Fig. 6G and Supplementary Fig. 5A), ACE2-A549 cells (Supplementary Fig. 5B) and Vero E6 cells (Fig. 6H and Supplementary Fig. 5C) that were infected by S-GFP-LV during 4 hours in presence (or not) of OxATP, BzATP or serum of convalescent COVID-19 patients (Fig. 6, G, H and Supplementary Fig. 5, A-C) and revealed that purinergic receptor P2X7 does not regulate SARS-CoV-2 entry into host cells. Similarly, we analyzed the frequency of GFP^+^ cells detected after 48-hour infection of ACE2-TMPRSS2-A549 cells with S-GFP-LV in presence (or not) of Tranilast, YVAD or serum of convalescent patients. We observed that Tranilast and YVAD did not reduce the frequency of GFP^+^ cells, as compared to cells treated with serum of convalescent patients (Fig. 6, I-N). These results were confirmed by detecting the intracellular HIV-1 Cap24 capsid in ACE2-TMPRSS2-A549 cells (Fig. 6O and Supplementary Fig. 5D) and ACE2-A549 cells (Fig. 6P and Supplementary Fig. 5E) that were infected with S-GFP-LV during 4 hours in presence (or not) of Tranilast. Since ACE2 expression was shown to control viral entry (57), Vero E6 cells (Supplementary Fig. 6, A and B) and ACE2-A549 cells (Supplementary Fig. 6, C-F) were treated with OxATP, BzATP, Tranilast and/or YVAD, for 6 hours (Supplementary Fig. 6, A and B), 4 hours (Supplementary Fig. 6, C and D), or 24 hours (Supplementary Fig. 6, E and F), and ACE2 membrane expression was analyzed by flow cytometry. The expression of ACE2 did not change in the presence of the agonist of P2X7 (BzATP), the selective antagonist of P2X7 (OxATP), the selective inhibitor of NLRP3 (Tranilast) and the selective inhibitor of caspase-1 (YVAD), implying that the purinergic receptor P2X7, the protein NLRP3 and the caspase-1 do not affect the membrane expression of ACE2. Altogether, these results demonstrate that the activation of the purinergic receptor P2X7 and the NLRP3 inflammasome supports SARS-CoV-2 replication into target cells without modulating viral entry.

**Fig. 6.**
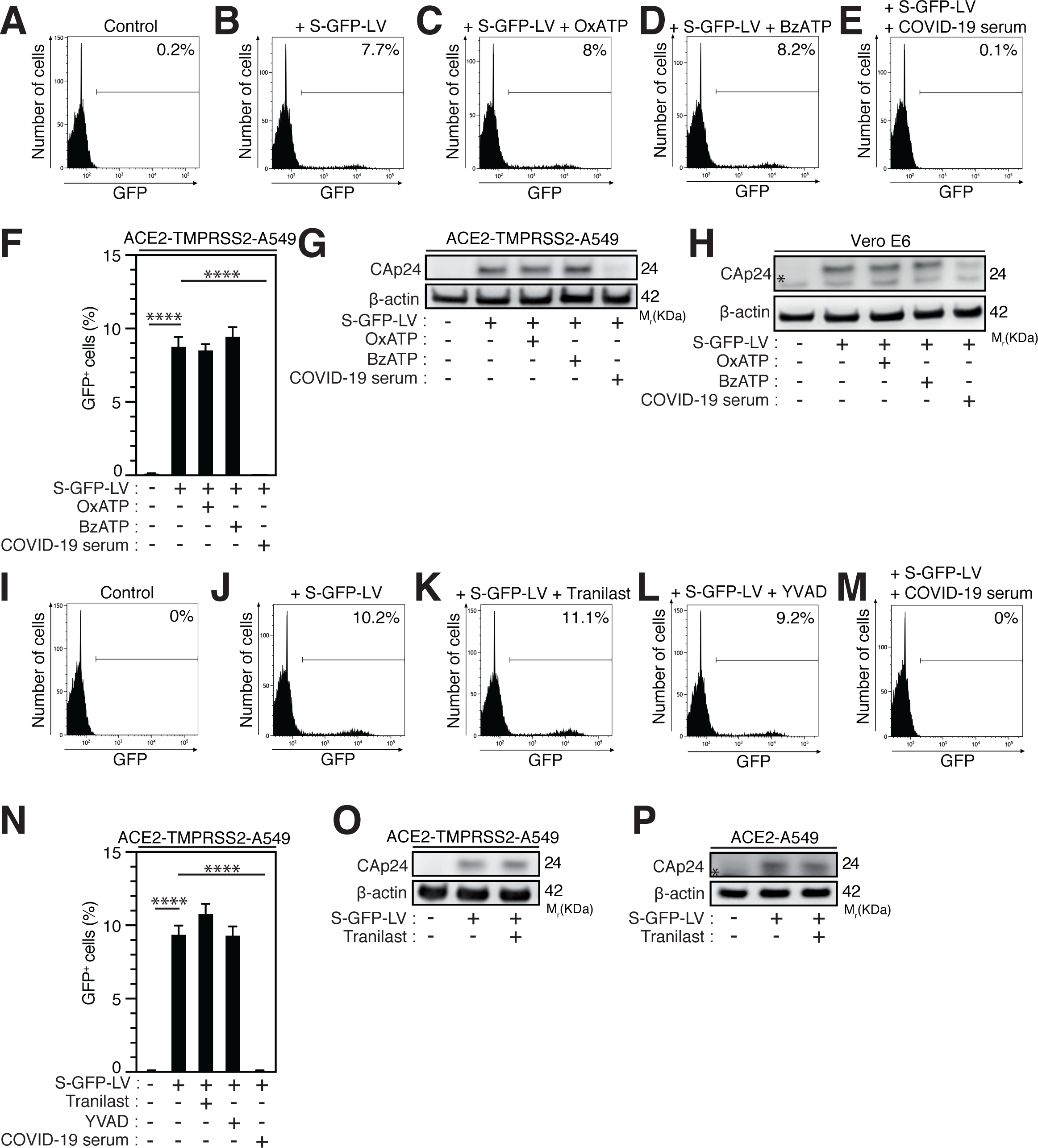
Purinergic receptor P2X7 and NLRP3 inflammasome control SARS-CoV-2 replication without affecting viral entry. **A-H** ACE2-TMPRSS2-A549 **(A-G)** or Vero E6 (**H)** cells were incubated during 4 hours with control (**A, B**), 100 µM OxATP (**C**), 100 µM BzATP (**D**), and infected for 48 hours (**B-F**) or 4 hours (**G, H**) with green fluorescent protein (GFP)-tagged HIV-1NL4-3**_Δ_**_Env_ variant (defective in viral envelope) pseudotyped with the SARS-CoV-2 spike (S) envelope (S-GFP-LV) or convalescent COVID-19 serum neutralized S-GFP-LV (**E-H**), and analyzed for GFP fluorescence by flow cytometry (**A-F**) or for intracellular HIV-1 CAp24 capsid and β-actin expression (**G, H**). Representative flow cytometry analysis (**A-E**) and percentages of GFP^+^ cells (**F**) are shown. **I-P** ACE2-TMPRSS2-A549 (**I-O**) or ACE2-A549 (**P**) cells were incubated during 4 hours with control (**I, J**), 100 µM Tranilast (**K**) or 100 µM YVAD (**L**) and infected for 48 hours (**J-L**) or 4 hours (**O, P**) with S-GFP-LV (**J-L, N-P**) or with convalescent COVID-19 serum neutralized S-GFP-LV (**M, N**). Then, cells were analyzed for GFP fluorescence by flow cytometry (**I-N**) or for intracellular HIV-1 CAp24 capsid and β-actin expression (**O, P**). Representative flow cytometry analysis (**I-M**) and percentages of GFP^+^ cells (**N**) are shown. In (**P**), the asterisk (*) is indicating a non-specific band. The data are presented as means ± SEM from at least 3 independent experiments. *p* values (\*\*\*\**p* < 0.0001) were determined using one-way ANOVA Tukey’s multiple comparisons test (**F, N**).

## Discussion

With the goal of studying the potential contribution of the purinergic receptor P2X7 and the NLRP3 inflammasome to the pathogenesis of SARS-CoV-2 infection, our study revealed that these two major sensors of danger signals participate to the early steps of SARS-CoV-2 infection. The activation of inflammasomes was detected during COVID-19 and positively correlates with disease severity in humans (29). Despite the fact that several inflammasomes were proposed to be activated and to contribute to organ dysfunction and disease severity during COVID-19 (29), the activation of NLRP3 inflammasome was recently associated with chronic stages of COVID-19 in humans (29) and in preclinical mouse models (30, 58). Several host targets (such as neutrophils, monocytes/macrophages and lymphocytes), which have been associated with the pathogenesis of SARS-CoV-2 infection (59), exhibited NLRP3 inflammasome activation during severe COVID-19. NLRP3-dependent inflammasome activation and uncontrolled extracellular traps (NET) production by neutrophils have been proposed to contribute to hyper-inflammation and dysregulated coagulation in COVID-19 patients with severe disease (60–62). Activation of the NLRP3 inflammasome and pyroptosis have been detected in monocytes and macrophages during SARS-CoV-2 infection (30, 31) and COVID-19 (29, 30, 63, 64). In addition, danger signals such as calprotectin recently associated with COVID-19 disease severity (65), are known to activate the NLRP3 inflammasome (66) or to be released as a consequence of its activation (67) and may contribute to pyroptosis (67). Accordingly, we detected the increased expression level of NLRP3 in alveolar macrophages and syncytium from autopsies of COVID-19 patients with severe disease, and observed the activation of NLRP3 inflammasome during the abortive infection of PMA-treated THP1 macrophages *in vitro*. Interestingly, our study also revealed that the NLRP3 inflammasome activation can also be detected in permissive host epithelial cells during early steps of SARS-CoV-2 infection and is required for SARS-CoV-2 replication. This process is rapidly induced after the interaction of SARS-CoV-2 with target cells, requires the activation of caspase-1 and acts positively at post-entry level on viral replication steps. Precise effects of NLRP3 inflammasome activation on cellular events (such as viral translation, polyprotein processing, double membrane vesicle (DMV) formation, replication organelle formation and activity) regulating SARS-CoV-2 replication (68) remain to be defined. In addition to its well-established role in triggering inflammation during microbial infection (53), our results highlighted an unexpected contribution of NLRP3 inflammasome activation to SARS-CoV-2 pathogenesis through the promotion of viral replication. Our results are in agreement with recent report revealing that pharmacological inhibition of NLRP3 inflammasome with MCC950 or knockout of NLRP3 abrogate both SARS-CoV-2 replication and associated inflammation during infection of human ACE2 mice (69). Considering the ability of the caspase-1 to regulate the activation of sterol regulatory element binding proteins (SREBPs), whose expressions have been detected increased in patients with severe COVID-19 (70) and genetic manipulations revealed their contributions to SARS-CoV-2 life cycle (68, 71, 72), further investigations to define the potential interplay between NLRP3 inflammasome activation, fatty acid and cholesterol metabolism and SARS-CoV-2 replication would be of interest and should help to identify key mechanisms regulating the replication of the SARS-CoV-2 that could be druggable targeted for COVID-19 treatment.

Our study demonstrated that the purinergic receptor P2X7 is also a key host factor in the SARS-CoV-2 replication cycle. The activation of purinergic receptor P2X7 supports SARS-CoV-2 replication without interfering with ACE2 membrane expression and viral entry. By deciphering the relationship between purinergic receptor P2X7 and NLRP3 inflammasome activation during the early steps of SARS-CoV-2 infection, our study demonstrated that purinergic receptor P2X7 acts upstream the activation of NLRP3 inflammasome. Since we previously observed that the purinergic receptors control HIV-1 envelope-elicited fusion (37, 39), it is conceivable that upon the interaction of spike (S) protein with host cell receptors and/or in response to the fusogenic activity of spike (S) protein, ATP is released from SARS-CoV-2 target cells and acts in an autocrine fashion on purinergic receptor P2X7 present at the membrane of SARS-CoV-2 target cells to stimulate NLRP3 inflammasome activation and to regulate viral replication.

The therapeutic inhibition of the key host factors of the signaling pathway that we identified in this study (P2X7èNLRP3èCaspase-1) could block the SARS-CoV-2 life cycle at the level of viral replication. Ivermectin, which is known to potentiate the activity of the purinergic receptor P2X4 (73), which can form heterotrimeric receptors with P2X7 (74), was recently found to inhibit the replication of SARS-CoV-2 in vitro (75), but did not shown benefits for COVID-19 patients (76). Nevertheless, further molecular investigations are required to fully understand the role of purinergic receptor family members and associated signaling pathways during early steps of SARS-CoV-2 infection and COVID-19-associated hyper-inflammation.

The clinical randomized clinical trials (RCT) having evaluating IL-1 inhibitors in severe COVID-19 yielded to controversial results. One RCT (NCT04362813) with canakinumab (a monoclonal antibody against anti-IL1β) was negative (77). With anakinra, an IL-1 receptor antagonist, one study (NCT04680949) in patients with increased soluble urokinase plasminogen receptor (suPAR) plasma level was positive (78), whereas 2 other trials (NCT04341584, NCT04364009) were negative (79). Our work suggests that targeting inflammasome-dependent IL-1β/IL-18 and/or purinergic signaling pathways may offer a novel opportunity for the treatment of viral infection and hyper-inflammation associated with COVID-19. Our study provides the first rationale for testing specific P2X7 antagonists such as CE-224,535 or JNJ-54175446, which were assessed without success for the treatment of rheumatoid arthritis (NCT00628095) and are under evaluation for the treatment of major depressive disorders (NCT04116606), respectively, alone or in combination with specific antagonists of NLRP3 such as Tranilast or Dapansutrile, which were assessed for the treatment of Cryopyrin-Associated Periodic Syndrome (CAPS) (NCT03923140) or systolic heart failures (NCT03534297), respectively. Repurposing of these existing drugs for blocking both viral replication and COVID-19-associated hyper-inflammation should rapidly improve the health of COVID-19 patients.

## Materials and methods

### Cell lines

The African green monkey kidney Vero E6 cells were purchased from ATCC (ATCC CRL-1587), ACE2-overexpressing A549 (ACE2-A549) cells were a gift from Dr. Olivier Schwartz (Institut Pasteur, France), ACE2 and transmembrane serine protease 2 (TMPRSS2)-expressing A549 (ACE2-TMPRSS2-A549) cells were from Invivogen (#a549-hace2tpsa) and HEK293T were from ATCC (#CRL-11268). These cells were cultured in Dulbecco’s modified Eagle’s medium (DMEM, Gibco, USA) with 10% heat inactivated fetal bovine serum (FBS), 100 UI/ml penicillin (Life technology), and 100 µg/ml streptomycin (Life technology) at 37°C. Caco-2 cells were kindly provided by Pr. Guido Kroemer (Gustave Roussy, France) and were cultured in EMEM (30-2003, ATCC) with 20% heat inactivated FBS, 100 UI/ml penicillin, and 100 µg/ml streptomycin at 37°C. Monocyte THP1 cells (TIB2002, ATCC) were differentiated in macrophages with phorbol-12 myristate-13-acetate (PMA) (#tlrl-pma, Invivogen), as previously described (80) and maintained in RMPI-1640-Glutamax medium supplemented with 10% heat inactivated FBS, 100 UI/ml penicillin, and 100 µg/ml streptomycin. All cell lines used were mycoplasma-free.

### Primary cells

Minor salivary gland biopsy specimens were obtained from patients referred for suspected primary Sjögren’s syndrome diagnosis to the rheumatology department of Bicêtre Hospital (AP-HP, Université Paris-Saclay). The subjects selected for this study were patients who presented with sicca symptoms but without any autoantibodies and with normal or subnormal minor salivary gland findings (i.e., focus score <1), and thus were considered as controls. Primary cultures of salivary gland epithelial cells were established from minor salivary glands based on previously described protocol (81). Briefly, the minor salivary gland biopsy was cut into small fragments, which were placed in 75-cm² flasks (BD Falcon) (prehydrated with 3 ml of s-BEM medium during 30 minutes at 37°C) with 1 ml of s-BEM medium. The supplemented-Basal Epithelial Medium (s-BEM) consisted of 3:1 mix of Ham’s F12 (Gibco, Life Technologies, UK) and DMEM (Gibco, Life Technologies, UK), supplemented with 2,5% FBS (PAN BIOTECH, Aidenbach, Germany), 10 ng/ml epidermal growth factor (EGF) (Gibco), 0.4 µg/ml hydrocortisone (Upjohn 100mg SERB), 0.5 µg/ml insulin (Novo Nordisk, Novorapid Flexpen) and 1× penicillin/streptomycin (LONZA, Verviers, Belgium). The flasks were incubated at 37°C with 5% CO2 in a humidified incubator. The day after, 3ml of s-BEM were added slowly in the flasks. At day 6, 5 ml of s-BEM were added. Then, cultures were fed with fresh medium twice a week. After biopsy, 2-3 small pieces were seeded into T75 flask and cultured in a DMEM-F12 medium supplemented with 2.5% FBS, 100 UI/ml penicillin/streptomycin, 0.4 µg/ml hydrocortisone, 0.5 µg/ml insulin and 10 ng/ml EGF. Once tissues had expended, cells were harvested and 1×10^5^ cells per well were plated in 48-well plates that were pretreated with collagen for 1 hour at 37°C. Twenty-four hours later, cells were treated with OxATP and infected with SARS-CoV-2 at a multiplicity of infection (MOI) of 0.5 for 48 hours. Cellular RNA was collected and extracted with NucleoSpin RNA Plus XS, Micro kit for RNA purification with DNA removal column (#740990.250, Macherey-Nagel), according to manufacturer instructions. Samples were then subjected to reverse transcription and PCR using probes against RdRp and β-actin. Data were normalized to β-actin and to control condition following the 2^-ΔΔCt^ method.

### Human autopsies and patients’ serum

Post-mortem lung sections were obtained from 3 non-COVID-19 patients and 7 COVID-19 patients with severe disease. Controls (n=3) were patients deceased after hemorragia, cardiorespiratory failure or interstitial pneumonia associated with pulmonary capillaritis. All COVID-19 patients (n=7) deceased after cardiorespiratory failure and exhibited in their vast majority diffuse alveolar damage (n=6) or fibrosis (n=1). Detection of SARS-CoV-2 was performed by RT-PCR on all patients using nasopharyngeal swabs. Convalescent COVID-19 serum was obtained from patients that received three doses of COVID-19 mRNA vaccine (Pfizer). Serum was collected one month after the last injection. All patients had a titer > 40 000 units/ml. The serum was obtained from Dr. Samuel Lebourgeois and Dr. Nadhira Houhou-Fidouh (Hôpital Bichat Claude Bernard, France).

### Virus, pseudoviral constructs and viral production

The SARS-CoV-2 BetaCoV/France/IDF0372/2020 strain (D614G) was provided by Dr. Benoit Visseaux from the group of Prof. Diane Descamps (UMR S 1135, Hôpital Bichat Claude Bernard, Paris) and by the National Reference Center For Respiratory Viruses (Institut Pasteur, Paris, France). The SARS-CoV-2 Alpha variant (lineage B1.1.7) was obtained form Pr. Mauro Pistello. Viral stocks were prepared by propagation in African green monkey kidney epithelial (Vero E6) cells in a biosafety level-3 (BLS-3) laboratory and titrated using lysis plaque assay as previously described (82). SARS-CoV-2 stock titer was 2×10^6^ PFU/ml. After 72 hours of infection, the supernatant was centrifuged at 1500 rpm, 12°C, for 5 minutes to remove dead cells and then centrifuged at 3000 rpm for 20 minutes at 12°C. Supernatant was then aliquoted and stored at −80°C. For the production of green fluorescent protein (GFP)-tagged HIV-1NL4-3**_Δ_**_env_ variant (defective in viral envelope) pseudotyped with the SARS-CoV-2 spike (S) envelope (S-GFP-LV), the plasmid NLENG1-ES-IRES coding for defective virus carrying two stop codons in the reading frame of the envelope and expressing the fluorescent reporter gene EGFP (GFP) was derived from the HIV-1 virus NL4-3 as previously published (83). Envelope defective virus was pseudotyped with HU-1 SARS-CoV-2 Spike protein (pLV-Spike, Invivogen) and used for single round infection assays. To produce viral stocks, 293T cells were transfected with virus-encoding plasmids by the calcium phosphate method. Briefly, 293T cells were transfected using 20 µg of NLENG1-ES-IRES and 4 µg of the corresponding plasmid encoding spike (S) envelop. Two days post-transfection, supernatants containing the viruses were submitted to a low-speed centrifugation step, filtered through a 0.45-µm low-protein-binding Durapore filter (Millipore), and stored at −80°C until use. Viral stocks were titrated using HIV-1 P24 ELISA assay (#NEK05001KT, Perkin Elmer) and the concentration was 200 ng/ml.

### Viral infections

Vero E6 cells, ACE2-TMPRSS2-A549, ACE2-A549 and Caco-2 cells were infected during indicated times with SARS-CoV-2 (BetaCoV/France/IDF0372/2020) at a MOI from 0.1 to 0.2 for qPCR experiments, MOI of 0.1 for WB experiments, and from MOI 1 to 2 for cytopathogenic analyses, in absence or in presence of 50 µM pyridoxal-phosphate-6-azopheny-2’,4’-disulfonate (PPADS) (#P178, Sigma-Aldrich), 100 µM oxidized ATP (OxATP) (#A6779, Sigma-Aldrich), 100 µM 2’(3’)-O-(4-Benzoylbenzoyl) adenosine 5’-triphosphate (BzATP) (#B6396, Sigma-Aldrich), 20 µM MCC950 (#inh-mcc, InvivoGen), 100 µM Tranilast (S-1439, DIVBIOSCIENCE) or 100 µM Ac-YVAD-cmk (#inh-yvad, InvivoGen), unless stated otherwise. The virus was also neutralized by mixing 1 µL of convalescent COVID-19 serum with 30 µl of virus for 40 minutes at room temperature, before added to the cells and incubated for 48 hours at 37°C. As previously described, primary salivary gland epithelial cells were obtained and infected in 48-wells and infected with SARS-CoV-2 at MOI 0.5 for 48 hours. For viral yielding experiment, Huh7 cells were inoculated with SARS-CoV-2 Alpha variant (lineage B1.1.7) at MOI 0.001 for 3 hours. Then, the viral inoculum was removed and cells were gently washed with ice-cold phosphate buffer saline (PBS). Fresh cell culture medium was added and cells were incubated for 24 hours. Supernatants were then collected and the viral RNA was extracted by automated RNA extractor EZ1 (Quiagen) and quantified using quantitative RT-PCR.

### RNA interference

The small interfering RNAs (siRNAs) against NLRP3 (siNLRP3) (5’-UGCAAGAUCUCUCAGCAAA-3’) and the corresponding control siRNAs (siControl) (5’-UUCAAUAAAUUCUUGAGGU-5’) were synthesized from Sigma. The siGenome smart pool siRNAs and siON Target plus siRNAs were purchased from Dharmacon and are composed of a pool of four siRNAs. Sequences of indicated siRNAs are following: siGenome smart pool siRNAs against CASP-1 (siCASP-1, #M-004401-03-0010): (1) 5’-GACUCAUUGAACAUAUGCA-3’, (2) 5’-AGACAUCCCACAAUGGGCU-3’, (3) 5’-GAAUAUGCCUGUUCCUGUG-3’, (4) 5’-CCGCAAGGUUCGAUUUUCA-3’, siGenome smart pool non-targeting siRNAs (siCo., #D-001206-14-05): (1) 5’-UAAGGCUAUGAAGAGAUAC-3’, (2) 5’-AUGUAUUGGCCUGUAUUAG-3’, (3) 5’-AUGAACGUGAAUUGCUCAA-3’, (4) 5’-UGGUUUACAUGUCGACUAA-3’, siON-Target plus siRNAs against P2X7 (siP2X7, #L-003728-00-0005): (1) 5’-GCGGUUGUGUCCCGAGUAU-3’, (2) 5’-GGAUCCAGAGCAUGAAUUA-3’, (3) 5’-GCUUUGCUCUGGUGAGUGA-3’, (4) 5’-GGAUAGCAGAGGUGAAAGA-3’, and the siON-Target plus non-targeting siRNAs (siCo., #D-001810-10-20): (1) 5’-UGGUUUACAUGUCGACUAA-3’, (2) 5’-UGGUUUACAUGUUGUGUGA-3’, (3) 5’UGGUUUACAUGUUUUCUGA-3’, (4) 5’UGGUUUACAUGUUUUCCUA-3’. Cells were seeded (10^5^ cells/well of 12 well-plate) and let be attached for 4 hours before siRNAs transfection. The siRNAs transfections were performed using lipofectamine RNAimax (#13778075, ThermoFisher Scientific) and OptiMEM (#31985-062, ThermoFisher Scientific), according to the manufacturer’s instructions. After 48 hours of transfection, cell were infected and analyzed for gene expression (by RT-PCR) or protein expression (by western blot).

### LentiCRISPR targeting

The generation of NLRP3-depleted THP1 cell line (CrNLRP3) was performed with two predesigned and validated CRISPR NLRP3 gRNAs (84) (gRNA1: CGAAGCAGCACTCATGCGAG; gRNA2: GTCTGATTCCGAAGTCACCG) which were cloned into the pLentiCRISPR v2 lentiviral vector expressing *CAS9* gene (GenScript). The control THP1 cell line (CrCo.) was obtained with the empty pLentiCRISPR v2 lentiviral vector. For the production of the lentiviral vectors (LentiCRISPR v2, LentiCRISPR v2-gRNA1-NLRP3 and LentiCRISPR v2-gRNA2-NLRP3), 10×10^6^ HEK293T (ATCC#CRL-3216) cells in a T300 plate and 20 ml DMEM medium (Gibco, #31966-021) (containing 2% heat inactivated FBS, 100 U/ml penicillin, 100 μg/ml streptomycin) were transfected with 8 µg pDM2-VSV-G, 20 µg pΔ8.91 packaging and 16 µg lentiviral vector plasmid using Fugene (Promega, #E2312) according manufacturer’s instructions. After 24 hours of transfection, cell supernatants were removed and replaced by 20 ml fresh DMEM medium (containing 2% heat inactivated FBS, 100 U/ml penicillin, and 100 μg/ml streptomycin) for additional 96 hours. The viral stocks in the cell supernatants were then harvested, filtered through a 0.45-µm-pore size filter and quantified for viral Cap24 content using enzyme-linked immunosorbent assay (ELISA) (Perkin Elmer, # NEK050A) as previously published (41, 85). CrCo. THP1 cells (1×10^6^) were transduced with 10 μg Cap24 LentiCRISPR v2 lentiviral vector and CrNLRP3 THP1 cells (1×10^6^) were transduced with 5 μg Cap24 of each lentiviral vector (LentiCRISPR v2-gRNA1-NLRP3 and LentiCRISPR v2-gRNA2-NLRP3). Transductions were performed in the presence of 5 µg/ml polybrene (Sigma, #H9268) by 2-hour spinoculation (1200 g at 25 °C) and 2-hour incubation at 37°C. Then, THP1 cells were washed and cultured for 72 hours after transduction before puromycin (InvivoGen, #ant-pr) selection (0.5 µg/ml) and validation of NLRP3 depletion by western blot analysis and qPCR.

### Immunofluorescence

Cells were fixed for 20 minutes with 4% paraformaldehyde-PBS, permeabilized with 0.5% Triton X-100 (#X100, Sigma-Aldrich) for 15 minutes and incubated with 5% BSA (Bovine Serum Albumin) in PBS for 1 hour at room temperature. After saturation, cells were incubated overnight at 4°C with anti-TMS1/ASC antibody (ASC) (#ab155970 rabbit, Abcam, 1/200 dilution) or anti-spike (S) antibody (#GTX632604 mouse, Genetex, 1/100 dilution) in PBS-BSA 5%. Then, cells were incubated with appropriate secondary antibody conjugated to Alexa Fluor 488 (#A11034, Invitrogen, 1/500) or Alexa Fluor 546 (#A11030, Invitrogen, 1/500) at room temperature for 1 hour. Nuclei were stained with Hoechst 33342 (#1874027, Invitrogen, 1/1000 dilution) for 30 minutes at room temperature. After three PBS washes, cells were mounted with Fluoromount G medium (Southern biotech). Cells were analyzed and imaged by Leica Dmi8 microscope using a 60X objective.

### Western blots

Total cellular proteins were extracted in a lysis buffer containing 1% NP-40, 0.5 M EDTA, 20 mM HEPES, 10 mM KCl, 1% glycerol and the protease and phosphatase inhibitors (Roche). 10-40 µg of protein extracts were run on 4–12% SDS-PAGE and transferred at 4°C onto a nitrocellulose membrane (0.2 Micron). After 1 hour saturation at room temperature with 5% BSA in Tris-buffered saline and 0.1% Tween 20 (TBS-Tween), nitrocellulose membranes were incubated with primary antibody at 4°C overnight. Horseradish peroxidase (HRP)-conjugated goat anti–mouse or anti–rabbit (SouthernBiotech) antibodies were then incubated for 1 hour and revealed with the enhanced ECL detection system (GE Healthcare). The primary antibody against P2X7 (#APR-004) was obtained from Alomone laboratories. Primary antibodies against ACE2 (#AF933) and Spike (S) (#GTX632604) were from R&D Systems and Genetex, respectively. Primary antibody against β-Actin-HRP (#ab49900) was purchased from Abcam. Anti-NLRP3 (Cryo-2, #AG-20B-0014) was from Adipogen. P24 antibody was obtained through the NIH HIV Reagent Program (Division of AIDS, NIAID, NIH: Polyclonal Anti-Human Immunodeficiency Virus Type 1 SF2 p24 (antiserum, Rabbit), ARP-4250, contributed by DAIDS/NIAID; produced by BioMolecular Technologies).

### Flow cytometry

Control, treated-ACE2-A549 and treated-Vero E6 cells were harvested in DMEM medium, washed three times in PBS and saturated with 2% PBS-BSA at 4°C for 20 minutes. Primary anti-human ACE-2 antibody (#AF933, R&D systems) or isotype-matched antibody (#AB-108-C, R&D systems) was incubated at 0.2 µg/10^6^ cells for ACE2-A549 and at 0.4 µg/10^6^ cells for Vero E6 cells during 30 minutes at room temperature in 2% PBS-BSA. Cells were washed three times with 2% PBS-BSA and incubated with secondary anti-goat IgG (H+L) APC-conjugated antibody (F0108, R&D systems) in 2% PBS-BSA for 20 minutes at room temperature at 0.05 µg/10^6^ cells for ACE2-A549 cells or at 0.1 µg/10^6^ cells for Vero E6 cells. Cells were then washed three times with 2% PBS-BSA, resuspended in 200 µl 2% PBS-BSA and analyzed for mean fluorescence intensity (MFI) of ACE2 expression with CytoFLEX flow cytometer (Beckman). To determine the biological effects of the purinergic receptor P2X7 and NLRP3 inflammasome on viral entry, ACE2-TMPRSS2-A549 cells were pretreated for 4 hours with indicated drugs and infected for 24 hours with SPIKE-GFP-LVs. As negative control, spike-GFP-LVs were preincubated with convalescent COVID19 patient’s serum during 40 minutes at room temperature. After extensive washes, ACE2-TMPRSS2-A549 cells were incubated in complete medium for 24 hours, fixed with 4% PFA and GFP-positive cells were analyzed using BD FACSCelesta flow cytometer (BD Biosciences).

### Quantitative RT-PCR

After infection, total cell RNA was extracted using either QIAshredder kit (#79654, Qiagen) and the RNeasy kit plus Mini kit (#74134, Qiagen) or using NucleoSpin RNA Plus XS, Micro kit for RNA purification with DNA removal column (#740990.250, Macherey-Nagel), according to the manufacturer’s instructions. RNA was first submitted to reverse transcription before PCR amplification. Quantifications were performed by real-time PCR on a Light Cycler instrument (Roche Diagnostics, Meylan, France) using the second-derivative-maximum method provided by the Light Cycler quantification software (version 3.5 (Roche Diagnostics)) or using CFX Maestro (BioRad). Standard curves for SARS-CoV-2 RNA quantifications were provided in the quantification kit and were generated by amplification of serial dilutions of the provided positive control. Sequences of the oligonucleotides and probes used to quantify the RdRp (F: 5’-GGTAACTGGTATGATTT-3’; R: 5’-CTGGTCAAGGTTAATATA-3’; Probe: 5’-[FAM]TCATACAAACCACGCCAGG[BHQ1]-3’), E gene (F: 5’-ACAGGTACGTTAATAGTTAATAGCGT-3’; R: 5’-ATATTGCAGCAGTACGCACACA-3’; Probe: 5’-[FAM]CAGGTGGAACCTCATCAGGAGATGC[BBQ]-3’) and β-actin (F 5’-CACCATTGGCAATGAGCGGTTC-3’; R 5’-AGGTCTTTGCGGATGTCCACGT-3’; SYBR-Green) are respectively obtained from Eurofins, TIB MolBiol or Origene (NM_001101). The amplification of RdRp and E genes was obtained after 5 minutes of reverse transcription, 5 minutes of denaturation and 45 to 50 cycles with the following steps (95°C during 5 seconds, 60°C during 15 seconds and 72°C during 15 seconds or 95°C for 15 seconds and 58°C for 30 seconds to 1 minute). The amplification of CASP-1 (Catalog #4331182 Assay ID Hs00354836_m1, ThermoFisher Scientific) and NLRP3 (Catalog #4331182 Assay ID Hs00918082_m1, ThermoFisher Scientific) was obtained after 5 minutes at 65°C for annealing primer to the template RNA, 2 minutes at 50°C of reverse transcription, 10 minutes at 95°C of denaturation and 44 cycles with the following steps (95°C for 15 seconds and 60°C for 1 min). Cq results were normalized by the total amount of RNA in the sample or by β-actin mRNA expression levels and reported to the condition without any compound (control condition). Data are presented as fold changes and were calculated with relative quantification of ΔCT obtained from quantitative RT-PCR. For the detection of SARS-CoV-2 in the supernatant of infected Huh7 cells, the viral RNA was extracted by automated RNA extractor EZ1 (Quiagen) and real-time RT-PCR was performed by targeting the SARS-CoV-2 RdRp gene as follows: each 20 µl sample consisted of 12.5 µl One Step PrimeScript™ III RT-PCR Kit (Takara Bio, Shiga, Japan), 0.5 µM Sars-CoV-2 CRV forward primer (5’-TCACCTAATTTAGCATGGCCTCT-3’), 0.5 µM SARS-CoV-2 CRV reverse primer (5’-CGTAGTGCAACAGGACTAAGC-3’), 0.1 µM SARS-CoV-2 CRV probe (5’-FAM-ACAGCAGAATTGGCCCTTAAAGCT-BHQ1-3’), 4 µl of purified nucleic acid and nuclease-free water. The in-house one-step RT-PCR reaction mixtures were run in a CFX Connect Real-Time PCR instrument (Bio-Rad Laboratories, Hercules, CA, USA) using previously standardized thermal conditions (52°C for 5 minutes, 95°C for 10 seconds, 45 cycles of 10 seconds at 95°C, and 62°C for 30 seconds).

### Detection of IL-1β cytokine release

Supernatants harvested from PMA-stimulated THP1 macrophages, Caco-2 cells or ACE2-A549 cells that were treated with MCC950, depleted for NLRP3 or incubated with COVID-19 patient’s serum, and infected as indicated with SARS-CoV-2, or from control uninfected cells were analyzed using ELISA assay for IL-1β, according to the manufacturers’ instructions. IL-1β secretion from PMA-THP1 was detected using IL-1 beta Human ELISA kit (#KHC0011, ThermoFisher Scientific) and IL-1 beta Human ELISA kit, High Sensitivity (#BMS224HS, ThermoFisher Scientific) was used for the detection of IL-1β secretion from epithelial cells.

### Cell viability assays

Vero E6 cells and ACE2-A549 cells were pretreated with indicated concentrations of PPADS, OxATP and/or BzATP during 4 hours before infection and infected or not with SARS-CoV-2 BetaCoV/France/IDF0372/2020 strain with a multiplicity of infection between 1 and 2. Cell viability was determined after 72 hours using 3-(4,5-dimethylthiazol-2-yl)-2,5-diphenyltetrazolium bromide (MTT) assay (#M5655, Sigma) following manufacturer’s instructions. Cell viability was also performed in uninfected Vero E6 cells, Caco-2 cells, ACE2-A549 cells and primary salivary gland epithelial cells with the same compounds incubated for 48 hours.

### Immunohistochemistry

Post-mortem lung specimens were fixed in formalin and embedded in paraffin. Tissue sections were deparaffinized, rehydrated, incubated in 10 mM sodium citrate, pH 6.0, microwaved for antigen retrieval and treated with 3% H_2_O_2_ to block endogenous peroxidase activity. Then, mouse antibodies against NLRP3/NALP3 (#AG-20B-0014, AdipoGen) (1:100), CD68 (#KP-1, Ventana) (prediluted), or double-stranded RNA (#J2-2004, Scicons J2) (1:500) and biotinylated goat anti-mouse IgG (#BA-9200, Vector) were incubated with lung sections. Immuno-reactivities were visualized using avidin-biotin complex-based peroxidase system (#PK-7100, Vector) and 3,3’-diaminobenzidine (DAB) peroxidase (HRP) substrate Kit (#SK-4100, Vector). Lung sections were also stained with hematoxylin and eosin, as previously described (86) and assessed by two independent observers without the knowledge of clinical diagnosis, using a Leica DM2500 LED Optical microscope and a 63x objective.

### Statistical analysis

Statistical analysis was performed using GraphPad Prism 8.0 (GraphPad). Statistical tests and calculated *p* values (\**p*< 0.05, \*\**p* < 0.01, \*\*\**p*< 0.001, and \*\*\*\**p*< 0.0001) are indicated in each figure and corresponding figure legend.

## Author contribution

Jean-Luc Perfettini provided financial support. Jean-Luc Perfettini designed and conducted the study. Déborah Lécuyer, Roberta Nardacci, Désirée Tannous, Emie Gutierrez-Mateyron, Aurélia Deva-Nathan, Frédéric Subra, Cristina Di Primio, Paola Quaranta, Vanessa Petit, Clémence Richetta, Ali Mostefa-Kara, Franca Del Nonno, Laura Falasca, Romain Marlin, Pauline Maisonnasse, Julia Delahousse, Juliette Pascaud, Marie Naigeon, Olivier Delelis and Awatef Allouch performed experiments. Xavier Mariette provided cultures of salivary gland epithelial cells. Déborah Lécuyer, Roberta Nardacci, Désirée Tannous, Emie Gutierrez-Mateyron, Aurélia Deva-Nathan, Frédéric Subra, Cristina Di Primio, Paola Quaranta, Vanessa Petit, Ali Mostefa-Kara, Franca Del Nonno, Laura Falasca, Eric Deprez, Nathalie Chaput, Angelo Paci, Véronique Saada, David Ghez, Xavier Mariette, Mario Costa, Mauro Pistello, Awatef Allouch, Olivier Delelis, Mauro Piacentini, Roger Le Grand and Jean-Luc Perfettini analyzed the results. Déborah Lécuyer, Aurélia Deva-Nathan, Emie Gutierrez-Mateyron and Désirée Tannous assembled the figures. Déborah Lécuyer, Awatef Allouch, Frédéric Subra and Jean-Luc Perfettini wrote the initial draft. Roberta Nardacci, Cristina Di Primio, Paola Quaranta, Vanessa Petit, Eric Deprez, Nathalie Chaput, Angelo Paci, Véronique Saada, David Ghez, Xavier Mariette, Mario Costa, Mauro Pistello, Olivier Delelis, Mauro Piacentini and Roger Le Grand provided advice and edited the initial draft.

## Ethics

This non-interventional study received approval by the institutional review board of the National Institute for Infectious Disease « Lazzaro Spallanzani » and following the principles stated in the Declaration of Helsinski. Human autopsies were performed at the National Institute for Infectious Diseases Lazzaro Spallanzani-IRCCS Hospital (Rome, Italy) according to guidance for post-mortem collection and submission of specimens and biosafety practices (CDC March 2020, Interim Guidance and (87)) to reduce the risk of transmission of infectious pathogens during and after the post-mortem examination. Autopsies were performed in accordance with the law owing to the unknown cause of death, and to both scientific and public interest in a pandemic novel disease. No informed consent was obtained from the families. All performed procedures and investigations were in accordance with the ethical standards of the Institutional Review Board of Lazzaro Spallanzani National Institute for Infectious Disease (Ethics Committee Approval number: n° 9/2020).

## Availability of Data and Materials

The datasets generated and/or analyzed during the current study are available from the corresponding author on reasonable request.

## Supporting information

LECUYER bioRxiv 2023 Supplementary Materials

## Acknowledgements

We gratefully acknowledge NH TherAguix and ENS Paris-Saclay for supporting our effort against COVID-19 pandemic, Dr. F. Griscelli and Dr. E. Gallois from Biology and Pathology department at Gustave Roussy hospital, Y. Lecluse and S. Salome-Desnoulez from Gustave Roussy’s imaging and cytometry platform (PFIC) for their technical support. We thank the staff of the animal facility of IDMIT, particularly B. Delache, M. Pottier, S. Langlois, JM. Robert and F. Relouzat.

## References

1. Baden LR, El Sahly HM, Essink B, Kotloff K, Frey S, Novak R, et al. Efficacy and Safety of the mRNA-1273 SARS-CoV-2 Vaccine. N Engl J Med. 2021;384(5):403–16.

2. Polack FP, Thomas SJ, Kitchin N, Absalon J, Gurtman A, Lockhart S, et al. Safety and Efficacy of the BNT162b2 mRNA Covid-19 Vaccine. N Engl J Med. 2020;383(27):2603–15.

3. Sadoff J, Gray G, Vandebosch A, Cardenas V, Shukarev G, Grinsztejn B, et al. Safety and Efficacy of Single-Dose Ad26.COV2.S Vaccine against Covid-19. N Engl J Med. 2021;384(23):2187-201.

4. Chen P, Nirula A, Heller B, Gottlieb RL, Boscia J, Morris J, et al. SARS-CoV-2 Neutralizing Antibody LY-CoV555 in Outpatients with Covid-19. N Engl J Med. 2021;384(3):229–37.

5. Gupta A, Gonzalez-Rojas Y, Juarez E, Crespo Casal M, Moya J, Falci DR, et al. Early Treatment for Covid-19 with SARS-CoV-2 Neutralizing Antibody Sotrovimab. N Engl J Med. 2021;385(21):1941–50.

6. Weinreich DM, Sivapalasingam S, Norton T, Ali S, Gao H, Bhore R, et al. REGN-COV2, a Neutralizing Antibody Cocktail, in Outpatients with Covid-19. N Engl J Med. 2021;384(3):238–51.

7. Beigel JH, Tomashek KM, Dodd LE, Mehta AK, Zingman BS, Kalil AC, et al. Remdesivir for the Treatment of Covid-19 - Final Report. N Engl J Med. 2020;383(19):1813–26.

8. Wahl A, Gralinski LE, Johnson CE, Yao W, Kovarova M, Dinnon KH3rd, et al. SARS-CoV-2 infection is effectively treated and prevented by EIDD-2801. Nature. 2021;591(7850):451-7.

9. Jayk Bernal A, Gomes da Silva MM, Musungaie DB, Kovalchuk E, Gonzalez A, Delos Reyes V, et al. Molnupiravir for Oral Treatment of Covid-19 in Nonhospitalized Patients. N Engl J Med. 2022;386(6):509–20.

10. Hammond J, Leister-Tebbe H, Gardner A, Abreu P, Bao W, Wisemandle W, et al. Oral Nirmatrelvir for High-Risk, Nonhospitalized Adults with Covid-19. N Engl J Med. 2022;386(15):1397-408.

11. Hermine O, Mariette X, Tharaux PL, Resche-Rigon M, Porcher R, Ravaud P, et al. Effect of Tocilizumab vs Usual Care in Adults Hospitalized With COVID-19 and Moderate or Severe Pneumonia: A Randomized Clinical Trial. JAMA Intern Med. 2021;181(1):32–40.

12. Kalil AC, Patterson TF, Mehta AK, Tomashek KM, Wolfe CR, Ghazaryan V, et al. Baricitinib plus Remdesivir for Hospitalized Adults with Covid-19. N Engl J Med. 2021;384(9):795–807.

13. Heyer A, Günther T, Robitaille A, Lütgehetmann M, Addo MM, Jarczack D, et al. Remdesivir-induced emergence of SAR-CoV2 variants in patients with prolonged infection. Cell Reports Medicine. 2022.

14. Jackson CB, Farzan M, Chen B, Choe H. Mechanisms of SARS-CoV-2 entry into cells. Nat Rev Mol Cell Biol. 2022;23(1):3–20.

15. Hoffmann M, Kleine-Weber H, Schroeder S, Kruger N, Herrler T, Erichsen S, et al. SARS-CoV-2 Cell Entry Depends on ACE2 and TMPRSS2 and Is Blocked by a Clinically Proven Protease Inhibitor. Cell. 2020;181(2):271–80 e8.

16. Zang R, Gomez Castro MF, McCune BT, Zeng Q, Rothlauf PW, Sonnek NM, et al. TMPRSS2 and TMPRSS4 promote SARS-CoV-2 infection of human small intestinal enterocytes. Sci Immunol. 2020;5(47).

17. Matsuyama S, Nao N, Shirato K, Kawase M, Saito S, Takayama I, et al. Enhanced isolation of SARS-CoV-2 by TMPRSS2-expressing cells. Proc Natl Acad Sci U S A. 2020;117(13):7001–3.

18. Hillen HS, Kokic G, Farnung L, Dienemann C, Tegunov D, Cramer P. Structure of replicating SARS-CoV-2 polymerase. Nature. 2020;584(7819):154-6.

19. de Wit E, van Doremalen N, Falzarano D, Munster VJ. SARS and MERS: recent insights into emerging coronaviruses. Nat Rev Microbiol. 2016;14(8):523–34.

20. Bouhaddou M, Memon D, Meyer B, White KM, Rezelj VV, Correa Marrero M, et al. The Global Phosphorylation Landscape of SARS-CoV-2 Infection. Cell. 2020;182(3):685–712 e19.

21. Daniloski Z, Jordan TX, Wessels HH, Hoagland DA, Kasela S, Legut M, et al. Identification of Required Host Factors for SARS-CoV-2 Infection in Human Cells. Cell. 2021;184(1):92–105 e16.

22. Cao X. COVID-19: immunopathology and its implications for therapy. Nat Rev Immunol. 2020;20(5):269–70.

23. Lucas C, Wong P, Klein J, Castro TBR, Silva J, Sundaram M, et al. Longitudinal analyses reveal immunological misfiring in severe COVID-19. Nature. 2020;584(7821):463-9.

24. Han Y, Zhang H, Mu S, Wei W, Jin C, Tong C, et al. Lactate dehydrogenase, an independent risk factor of severe COVID-19 patients: a retrospective and observational study. Aging (Albany NY). 2020;12(12):11245–58.

25. Merad M, Blish CA, Sallusto F, Iwasaki A. The immunology and immunopathology of COVID-19. Science. 2022;375(6585):1122-7.

26. Cassel SL, Sutterwala FS. Sterile inflammatory responses mediated by the NLRP3 inflammasome. Eur J Immunol. 2010;40(3):607–11.

27. Swanson KV, Deng M, Ting JP. The NLRP3 inflammasome: molecular activation and regulation to therapeutics. Nat Rev Immunol. 2019;19(8):477–89.

28. Jorgensen I, Rayamajhi M, Miao EA. Programmed cell death as a defence against infection. Nat Rev Immunol. 2017;17(3):151–64.

29. Rodrigues TS, de Sa KSG, Ishimoto AY, Becerra A, Oliveira S, Almeida L, et al. Inflammasomes are activated in response to SARS-CoV-2 infection and are associated with COVID-19 severity in patients. J Exp Med. 2021;218(3).

30. Sefik E, Qu R, Junqueira C, Kaffe E, Mirza H, Zhao J, et al. Inflammasome activation in infected macrophages drives COVID-19 pathology. Nature. 2022;606(7914):585-93.

31. Ratajczak MZ, Bujko K, Ciechanowicz A, Sielatycka K, Cymer M, Marlicz W, et al. SARS-CoV-2 Entry Receptor ACE2 Is Expressed on Very Small CD45(-) Precursors of Hematopoietic and Endothelial Cells and in Response to Virus Spike Protein Activates the Nlrp3 Inflammasome. Stem Cell Rev Rep. 2020.

32. Shi CS, Nabar NR, Huang NN, Kehrl JH. SARS-Coronavirus Open Reading Frame-8b triggers intracellular stress pathways and activates NLRP3 inflammasomes. Cell Death Discov. 2019;5:101.

33. Chen IY, Moriyama M, Chang MF, Ichinohe T. Severe Acute Respiratory Syndrome Coronavirus Viroporin 3a Activates the NLRP3 Inflammasome. Front Microbiol. 2019;10:50.

34. Cekic C, Linden J. Purinergic regulation of the immune system. Nat Rev Immunol. 2016;16(3):177–92.

35. Burnstock G, Knight GE. Cellular distribution and functions of P2 receptor subtypes in different systems. Int Rev Cytol. 2004;240:31–304.

36. He Y, Franchi L, Nunez G. TLR agonists stimulate Nlrp3-dependent IL-1beta production independently of the purinergic P2X7 receptor in dendritic cells and in vivo. J Immunol. 2013;190(1):334–9.

37. Paoletti A, Allouch A, Caillet M, Saidi H, Subra F, Nardacci R, et al. HIV-1 Envelope Overcomes NLRP3-Mediated Inhibition of F-Actin Polymerization for Viral Entry. Cell Rep. 2019;28(13):3381–94 e7.

38. Di Virgilio F, Dal Ben D, Sarti AC, Giuliani AL, Falzoni S. The P2X7 Receptor in Infection and Inflammation. Immunity. 2017;47(1):15–31.

39. Seror C, Melki MT, Subra F, Raza SQ, Bras M, Saidi H, et al. Extracellular ATP acts on P2Y2 purinergic receptors to facilitate HIV-1 infection. J Exp Med. 2011;208(9):1823–34.

40. Paoletti A, Raza SQ, Voisin L, Law F, Caillet M, Martins I, et al. Editorial: Pannexin-1--the hidden gatekeeper for HIV-1. J Leukoc Biol. 2013;94(3):390-2.

41. Allouch A, Di Primio C, Paoletti A, Le-Bury G, Subra F, Quercioli V, et al. SUGT1 controls susceptibility to HIV-1 infection by stabilizing microtubule plus-ends. Cell Death Differ. 2020.

42. Ferrari D, Idzko M, Muller T, Manservigi R, Marconi P. Purinergic Signaling: A New Pharmacological Target Against Viruses? Trends Pharmacol Sci. 2018;39(11):926–36.

43. Carsana L, Sonzogni A, Nasr A, Rossi RS, Pellegrinelli A, Zerbi P, et al. Pulmonary post-mortem findings in a series of COVID-19 cases from northern Italy: a two-centre descriptive study. Lancet Infect Dis. 2020.

44. Ziegler CGK, Allon SJ, Nyquist SK, Mbano IM, Miao VN, Tzouanas CN, et al. SARS-CoV-2 Receptor ACE2 Is an Interferon-Stimulated Gene in Human Airway Epithelial Cells and Is Detected in Specific Cell Subsets across Tissues. Cell. 2020;181(5):1016–35 e19.

45. Chu H, Chan JF, Wang Y, Yuen TT, Chai Y, Hou Y, et al. Comparative Replication and Immune Activation Profiles of SARS-CoV-2 and SARS-CoV in Human Lungs: An Ex Vivo Study With Implications for the Pathogenesis of COVID-19. Clin Infect Dis. 2020;71(6):1400–9.

46. Merad M, Martin JC. Pathological inflammation in patients with COVID-19: a key role for monocytes and macrophages. Nat Rev Immunol. 2020;20(6):355–62.

47. Di Virgilio F. Liaisons dangereuses: P2X(7) and the inflammasome. Trends Pharmacol Sci. 2007;28(9):465–72.

48. Ahn JH, Kim J, Hong SP, Choi SY, Yang MJ, Ju YS, et al. Nasal ciliated cells are primary targets for SARS-CoV-2 replication in the early stage of COVID-19. J Clin Invest. 2021;131(13).

49. Hou YJ, Okuda K, Edwards CE, Martinez DR, Asakura T, Dinnon KH3rd, et al. SARS-CoV-2 Reverse Genetics Reveals a Variable Infection Gradient in the Respiratory Tract. Cell. 2020;182(2):429-46 e14.

50. Santana PT, Martel J, Lai HC, Perfettini JL, Kanellopoulos JM, Young JD, et al. Is the inflammasome relevant for epithelial cell function? Microbes Infect. 2016;18(2):93–101.

51. Martinon F, Burns K, Tschopp J. The inflammasome: a molecular platform triggering activation of inflammatory caspases and processing of proIL-beta. Mol Cell. 2002;10(2):417–26.

52. Ogura Y, Sutterwala FS, Flavell RA. The inflammasome: first line of the immune response to cell stress. Cell. 2006;126(4):659–62.

53. Franchi L, Munoz-Planillo R, Nunez G. Sensing and reacting to microbes through the inflammasomes. Nat Immunol. 2012;13(4):325–32.

54. Gurcel L, Abrami L, Girardin S, Tschopp J, van der Goot FG. Caspase-1 activation of lipid metabolic pathways in response to bacterial pore-forming toxins promotes cell survival. Cell. 2006;126(6):1135–45.

55. Paoletti A, Raza SQ, Voisin L, Law F, Pipoli da Fonseca J, Caillet M, et al. Multifaceted roles of purinergic receptors in viral infection. Microbes Infect. 2012;14(14):1278–83.

56. Matuck BF, Dolhnikoff M, Duarte-Neto AN, Maia G, Gomes SC, Sendyk DI, et al. Salivary glands are a target for SARS-CoV-2: a source for saliva contamination. J Pathol. 2021;254(3):239–43.

57. Glowacka I, Bertram S, Herzog P, Pfefferle S, Steffen I, Muench MO, et al. Differential downregulation of ACE2 by the spike proteins of severe acute respiratory syndrome coronavirus and human coronavirus NL63. J Virol. 2010;84(2):1198–205.

58. Sefik E, Israelow B, Mirza H, Zhao J, Qu R, Kaffe E, et al. A humanized mouse model of chronic COVID-19. Nat Biotechnol. 2022;40(6):906–20.

59. Lamers MM, Haagmans BL. SARS-CoV-2 pathogenesis. Nat Rev Microbiol. 2022;20(5):270–84.

60. Aymonnier K, Ng J, Fredenburgh LE, Zambrano-Vera K, Munzer P, Gutch S, et al. Inflammasome activation in neutrophils of patients with severe COVID-19. Blood Adv. 2022;6(7):2001–13.

61. Leppkes M, Knopf J, Naschberger E, Lindemann A, Singh J, Herrmann I, et al. Vascular occlusion by neutrophil extracellular traps in COVID-19. EBioMedicine. 2020;58:102925.

62. Veras FP, Pontelli MC, Silva CM, Toller-Kawahisa JE, de Lima M, Nascimento DC, et al. SARS-CoV-2-triggered neutrophil extracellular traps mediate COVID-19 pathology. J Exp Med. 2020;217(12).

63. Kroemer A, Khan K, Plassmeyer M, Alpan O, Haseeb MA, Gupta R, et al. Inflammasome activation and pyroptosis in lymphopenic liver patients with COVID-19. J Hepatol. 2020.

64. Junqueira C, Crespo A, Ranjbar S, de Lacerda LB, Lewandrowski M, Ingber J, et al. FcgammaR-mediated SARS-CoV-2 infection of monocytes activates inflammation. Nature. 2022;606(7914):576-84.

65. Silvin A, Chapuis N, Dunsmore G, Goubet AG, Dubuisson A, Derosa L, et al. Elevated Calprotectin and Abnormal Myeloid Cell Subsets Discriminate Severe from Mild COVID-19. Cell. 2020.

66. Simard JC, Cesaro A, Chapeton-Montes J, Tardif M, Antoine F, Girard D, et al. S100A8 and S100A9 induce cytokine expression and regulate the NLRP3 inflammasome via ROS-dependent activation of NF-kappaB(1.). PLoS One. 2013;8(8):e72138.

67. Basiorka AA, McGraw KL, Eksioglu EA, Chen X, Johnson J, Zhang L, et al. The NLRP3 inflammasome functions as a driver of the myelodysplastic syndrome phenotype. Blood. 2016;128(25):2960–75.

68. Baggen J, Vanstreels E, Jansen S, Daelemans D. Cellular host factors for SARS-CoV-2 infection. Nat Microbiol. 2021;6(10):1219–32.

69. Zeng J, Xie X, Feng XL, Xu L, Han JB, Yu D, et al. Specific inhibition of the NLRP3 inflammasome suppresses immune overactivation and alleviates COVID-19 like pathology in mice. EBioMedicine. 2022;75:103803.

70. Lee W, Ahn JH, Park HH, Kim HN, Kim H, Yoo Y, et al. COVID-19-activated SREBP2 disturbs cholesterol biosynthesis and leads to cytokine storm. Signal Transduct Target Ther. 2020;5(1):186.

71. Wang R, Simoneau CR, Kulsuptrakul J, Bouhaddou M, Travisano KA, Hayashi JM, et al. Genetic Screens Identify Host Factors for SARS-CoV-2 and Common Cold Coronaviruses. Cell. 2021;184(1):106–19 e14.

72. Schneider WM, Luna JM, Hoffmann HH, Sanchez-Rivera FJ, Leal AA, Ashbrook AW, et al. Genome-Scale Identification of SARS-CoV-2 and Pan-coronavirus Host Factor Networks. Cell. 2021;184(1):120–32 e14.

73. Stokes L, Layhadi JA, Bibic L, Dhuna K, Fountain SJ. P2X4 Receptor Function in the Nervous System and Current Breakthroughs in Pharmacology. Front Pharmacol. 2017;8:291.

74. Guo C, Masin M, Qureshi OS, Murrell-Lagnado RD. Evidence for functional P2X4/P2X7 heteromeric receptors. Mol Pharmacol. 2007;72(6):1447–56.

75. Caly L, Druce JD, Catton MG, Jans DA, Wagstaff KM. The FDA-approved drug ivermectin inhibits the replication of SARS-CoV-2 in vitro. Antiviral Res. 2020;178:104787.

76. Naggie S, Boulware DR, Lindsell CJ, Stewart TG, Gentile N, Collins S, et al. Effect of Ivermectin vs Placebo on Time to Sustained Recovery in Outpatients With Mild to Moderate COVID-19: A Randomized Clinical Trial. JAMA. 2022;328(16):1595–603.

77. Caricchio R, Abbate A, Gordeev I, Meng J, Hsue PY, Neogi T, et al. Effect of Canakinumab vs Placebo on Survival Without Invasive Mechanical Ventilation in Patients Hospitalized With Severe COVID-19: A Randomized Clinical Trial. JAMA. 2021;326(3):230–9.

78. Kyriazopoulou E, Poulakou G, Milionis H, Metallidis S, Adamis G, Tsiakos K, et al. Early treatment of COVID-19 with anakinra guided by soluble urokinase plasminogen receptor plasma levels: a double-blind, randomized controlled phase 3 trial. Nat Med. 2021;27(10):1752–60.

79. group C-C. Effect of anakinra versus usual care in adults in hospital with COVID-19 and mild-to-moderate pneumonia (CORIMUNO-ANA-1): a randomised controlled trial. Lancet Respir Med. 2021;9(3):295–304.

80. Wu Q, Allouch A, Paoletti A, Leteur C, Mirjolet C, Martins I, et al. NOX2-dependent ATM kinase activation dictates pro-inflammatory macrophage phenotype and improves effectiveness to radiation therapy. Cell Death Differ. 2017;24(9):1632–44.

81. Dimitriou ID, Kapsogeorgou EK, Abu-Helu RF, Moutsopoulos HM, Manoussakis MN. Establishment of a convenient system for the long-term culture and study of non-neoplastic human salivary gland epithelial cells. Eur J Oral Sci. 2002;110(1):21–30.

82. Gordon DE, Jang GM, Bouhaddou M, Xu J, Obernier K, White KM, et al. A SARS-CoV-2 protein interaction map reveals targets for drug repurposing. Nature. 2020;583(7816):459-68.

83. Levy DN, Aldrovandi GM, Kutsch O, Shaw GM. Dynamics of HIV-1 recombination in its natural target cells. Proc Natl Acad Sci U S A. 2004;101(12):4204–9.

84. Sanjana NE, Shalem O, Zhang F. Improved vectors and genome-wide libraries for CRISPR screening. Nat Methods. 2014;11(8):783–4.

85. Allouch A, Voisin L, Zhang Y, Raza SQ, Lecluse Y, Calvo J, et al. CDKN1A is a target for phagocytosis-mediated cellular immunotherapy in acute leukemia. Nat Commun. 2022;13(1):6739.

86. Falasca L, Nardacci R, Colombo D, Lalle E, Di Caro A, Nicastri E, et al. Post-Mortem Findings in Italian Patients with COVID-19 - a Descriptive Full Autopsy Study of cases with and without co-morbidities. J Infect Dis. 2020.

87. Hanley B, Lucas SB, Youd E, Swift B, Osborn M. Autopsy in suspected COVID-19 cases. J Clin Pathol. 2020;73(5):239–42.

