## Supplementary material for "The purinergic receptor P2X7 and the NLRP3 inflammasome are druggable host factors required for SARS-CoV-2 infection": LECUYER bioRxiv 2023 Supplementary Materials

**Supplementary Fig. 1 SARS-CoV-2 infection induces NLRP3 inflammasome activation in macrophages. A, B** The relative S expression data from Fig. 2A (**A**), Fig. 2B (**B**), Fig. 2G (**C**), Fig. 2H (**D**) and Fig. 2K (**E**) are presented as means ± SEM from at least 3 independent experiments. *p* values (***p* < 0.01 and ****p* < 0.001) were determined using one-way ANOVA Tukey's multiple comparisons test (**A-E**).

**Supplementary Fig. 2 MCC950 and Tranilast exhibited distinct effects on SARS-CoV-2 replication and did not affect cellular viability. A-C** ACE2-A549 cells were infected with SARS-CoV-2 at MOI = 2 for 48 hours with 20 μM MCC950 and evaluated for spike (S) and β-actin expressions by western bot (**A**, **B**) and spike (S) expression by fluorescence microscopy (**C**). Representative western blot (**A**), the relative S expression data from A (**B**) and representative image (**C**) are shown. Scale bar indicates 20 μm and DNA is detected using Hoechst 33342. **D, E** The relative S expression data from Fig. 3F (**D**) and Fig. 3G (**E**) are presented as means ± SEM from at least 3 independent experiments. **F-H** ACE2-A549 cells (**F, H**) or Caco-2 cells (**G**) were treated for 48 hours with 20 μM MCC950 (**F**) or indicated concentrations of Tranilast (**G, H**) and analyzed for viability using MTT assay. Percentages of viable cells are shown. Data are presented as means ± SEM from at least 3 independent experiments. *p* values (**p* < 0.05, ***p* < 0.01 and ****p* < 0.001) were determined using one-way ANOVA Tukey's multiple comparisons test (**D, G**) and unpaired t-test (**B**, **E**, **F** and **H**).

**Supplementary Fig. 3 Biological effects of YVAD and CASP-1 depletion on the viability of Caco-2 and ACE2-A549 cells or the relative expression of S protein. A, B** Caco-2 cells (**A**) or ACE2-A549 cells (**B**) were treated for 48 hours with 100 μM YVAD and analyzed for viability using MTT assay. Percentages of viable cells are shown. **C** The relative S expression data from Fig. 4H are presented as means ± SEM from at least 3 independent experiments. *p* values (**$p < 0.01$) were determined using unpaired t-test (**A-C**).

**Supplementary Fig. 4 Biological effects of OxATP, PPADS, BzATP or P2X7 depletion on the viability of Vero E6, ACE2-A549 and primary salivary gland epithelial cells or the relative expression of S protein.** **A-C** Vero E6 (**A, B**) and ACE2-A549 cells (**C**) were treated for 24 hours (**A, B**) or 48 hours (**C**) with indicated concentrations of OxATP (**A, C**) or PPADS (**B**) and analyzed for viability using MTT assay. Percentages of viable cells are shown. **D, E** The relative expression data for P2X7 (**D**) and S protein (**E**) from Fig. 5F are presented as means ± SEM from at least 3 independent experiments. **F** ACE2-A549 cells were treated for 48 hours with indicated concentrations of BzATP (**F**) and analyzed for viability using MTT assay. Percentages of viable cells are shown. **G** Primary salivary gland epithelial cells were treated for 48 hours with 100 μM of OxATP and analyzed for viability using MTT assay. Percentages of viable cells are shown. *p* values (*$p < 0.05$, **$p < 0.01$ and ***$p < 0.001$) were determined using one-way ANOVA Tukey's multiple comparisons test (**A-D** and **F**) and unpaired t-test (**E, G**).

**Supplementary Fig. 5 P2X7 modulation did not affect SARS-CoV-2 entry in ACE2-A549 cells.** **A** The relative CAp24 expression data from Fig. 6G are presented as means ± SEM from at least 3 independent experiments. **B** ACE2-A549 cells were treated with 100 μM of OxATP or BzATP and infected with S-GFP-LV or with convalescent COVID-19 serum neutralized S-GFP-LV and analyzed for intracellular HIV-1 CAp24 capsid and β-actin expression. A western blot representative of two independent experiments is shown. The asterisk (*) is indicating a non-specific band. **C** The relative CAp24 expression data from Fig.

6H are presented as means ± SEM from at least 3 independent experiments. **D** The relative CAp24 expression data from Fig. 6O are presented as means ± SEM from at least 3 independent experiments. **E** The relative CAp24 expression data from Fig. 6P are presented as means ± SEM from at least 3 independent experiments. *p* values (*$p < 0.05$, **$p < 0.01$ and ***$p < 0.001$) were determined using one-way ANOVA Tukey's multiple comparisons test (**A** and **C**) and unpaired t-test (**D, E**).

**Supplementary Fig. 6 Modulation of P2X7, NLRP3 inflammasome and caspase-1 biological activities did not affect membrane expression of ACE2. A-F** Vero E6 (**A, B**) or ACE2-A549 (**C-F**) cells were treated for 6 hours (**A, B**), 4 hours (**C, D**) or 24 hours (**E, F**) with 10 µM OxATP (**A**), 100 µM OxATP (**C, E**), 100 µM BzATP (**A, C** and **E**), 100 µM Tranilast or 100 µM YVAD (**B, D** and **F**) and analyzed for ACE2 membrane expression by flow cytometry. Fold change (FC) are shown. Data are presented as means ± SEM from at least 3 independent experiments. No statistically differences were found using one-way ANOVA Tukey's multiple comparisons test (**A-F**).

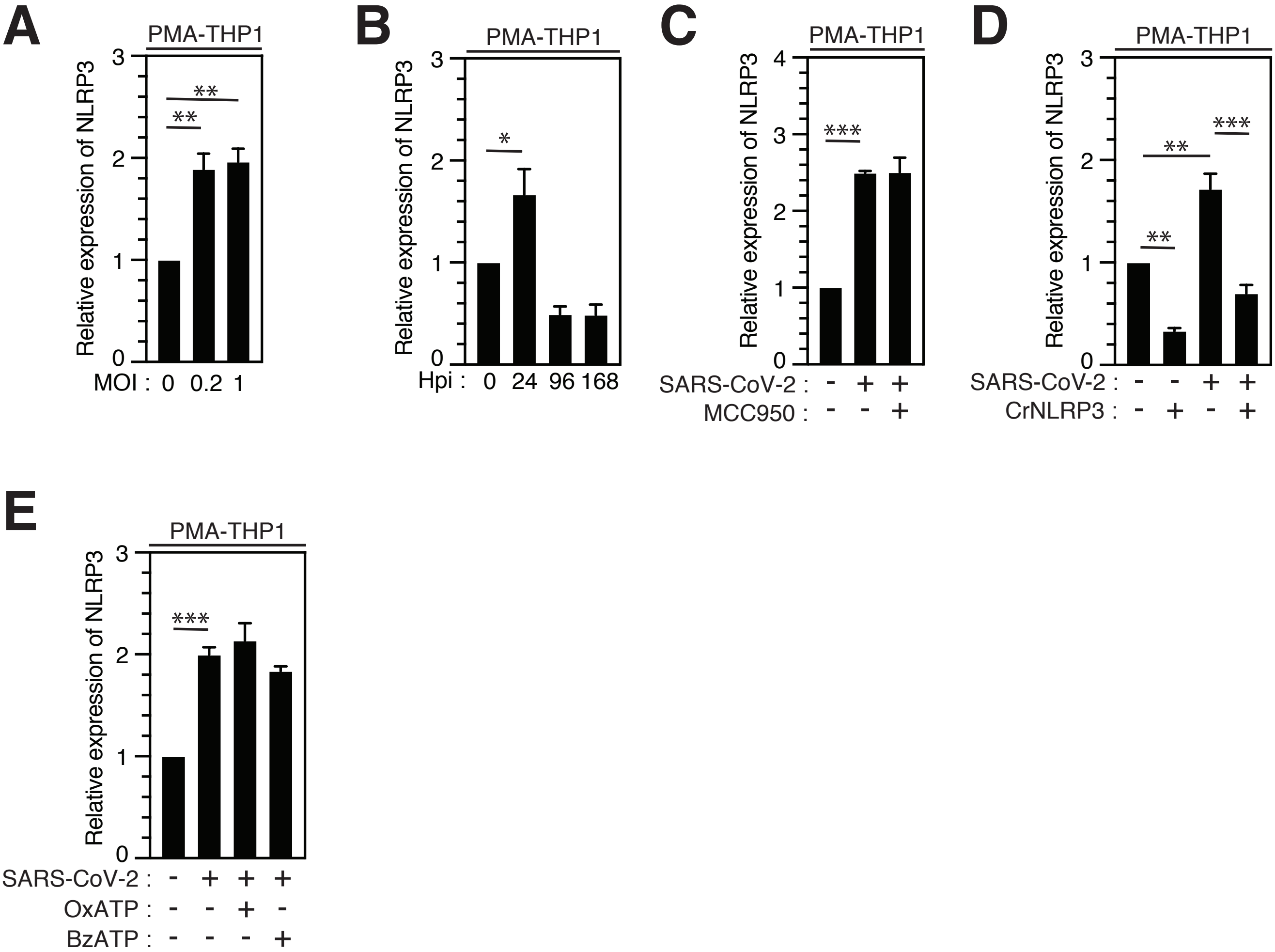
A
PMA-THP1
Relative expression of NLRP3
MOI : 0 0.2 1
B
PMA-THP1
Relative expression of NLRP3
Hpi : 0 24 96 168
C
PMA-THP1
Relative expression of NLRP3
SARS-CoV-2 : - + +
MCC950 : - - +
D
PMA-THP1
Relative expression of NLRP3
SARS-CoV-2 : - - + +
CrNLRP3 : - + - +
E
PMA-THP1
Relative expression of NLRP3
SARS-CoV-2 : - + + +
OxATP : - - + -
BzATP : - - - +

**LECUYER#S1**

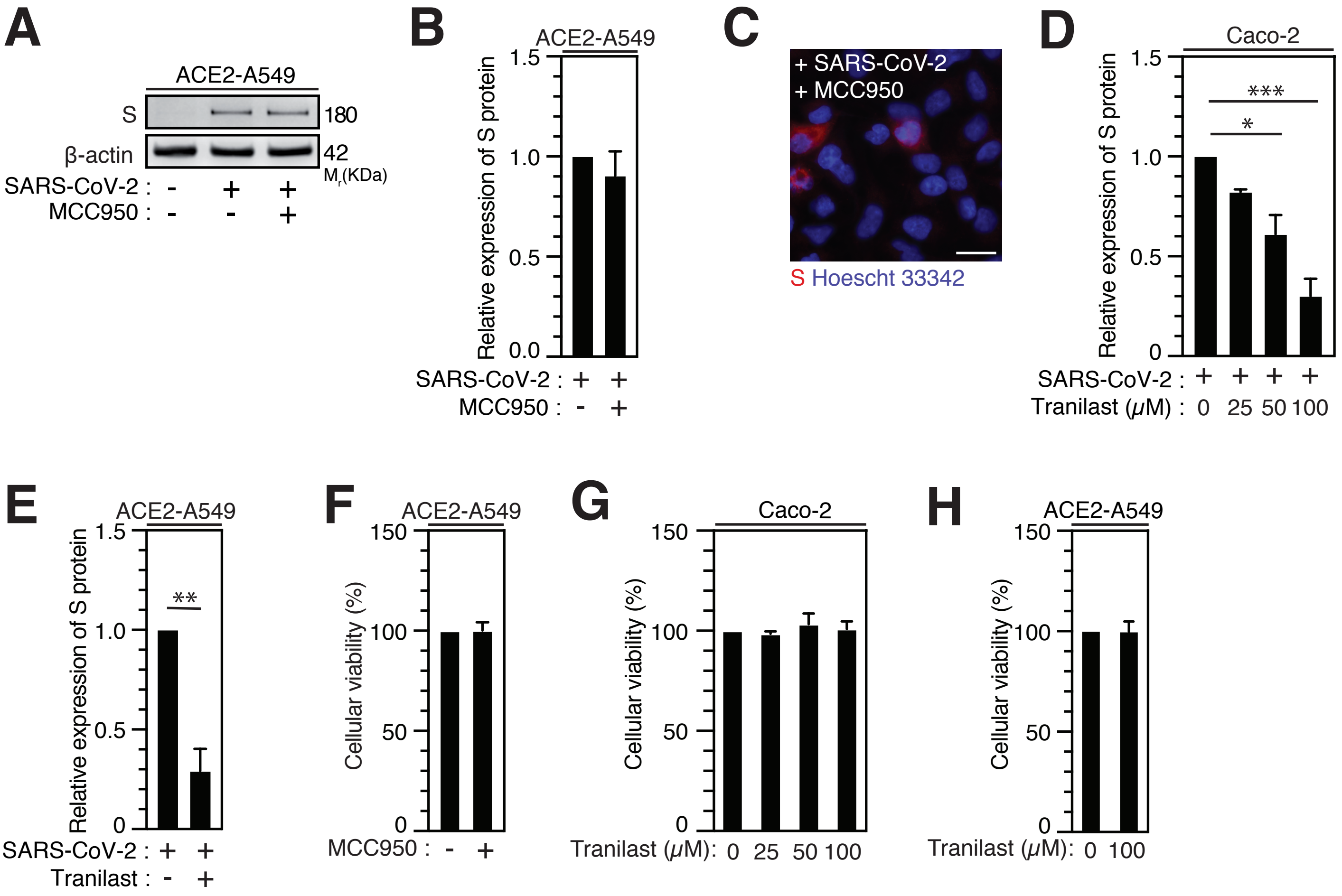
A
ACE2-A549
S
180
β-actin
42
Mr(KDa)
SARS-CoV-2 : - + +
MCC950 : - - +
B
ACE2-A549
Relative expression of S protein
SARS-CoV-2 : + +
MCC950 : - +
C
+ SARS-CoV-2
+ MCC950
S Hoescht 33342
D
Caco-2
Relative expression of S protein
SARS-CoV-2 : + + + +
Tranilast (μM) : 0 25 50 100
E
ACE2-A549
Relative expression of S protein
SARS-CoV-2 : + +
Tranilast : - +
F
ACE2-A549
Cellular viability (%)
MCC950 : - +
G
Caco-2
Cellular viability (%)
Tranilast (μM): 0 25 50 100
H
ACE2-A549
Cellular viability (%)
Tranilast (μM): 0 100

**LECUYER#S2**

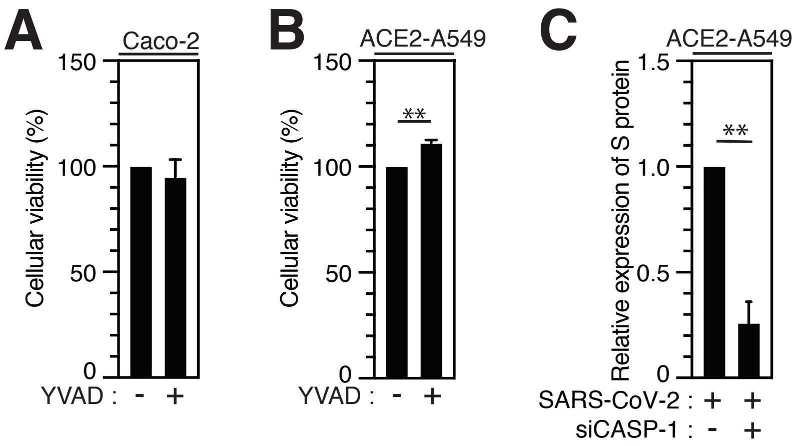
A
Caco-2
Cellular viability (%)
150
100
50
0
YVAD : - +
B
ACE2-A549
Cellular viability (%)
150
100
50
0
**
YVAD : - +
C
ACE2-A549
Relative expression of S protein
1.5
1.0
0.5
0
**
SARS-CoV-2 : + +
siCASP-1 : - +

**LECUYER#S3**

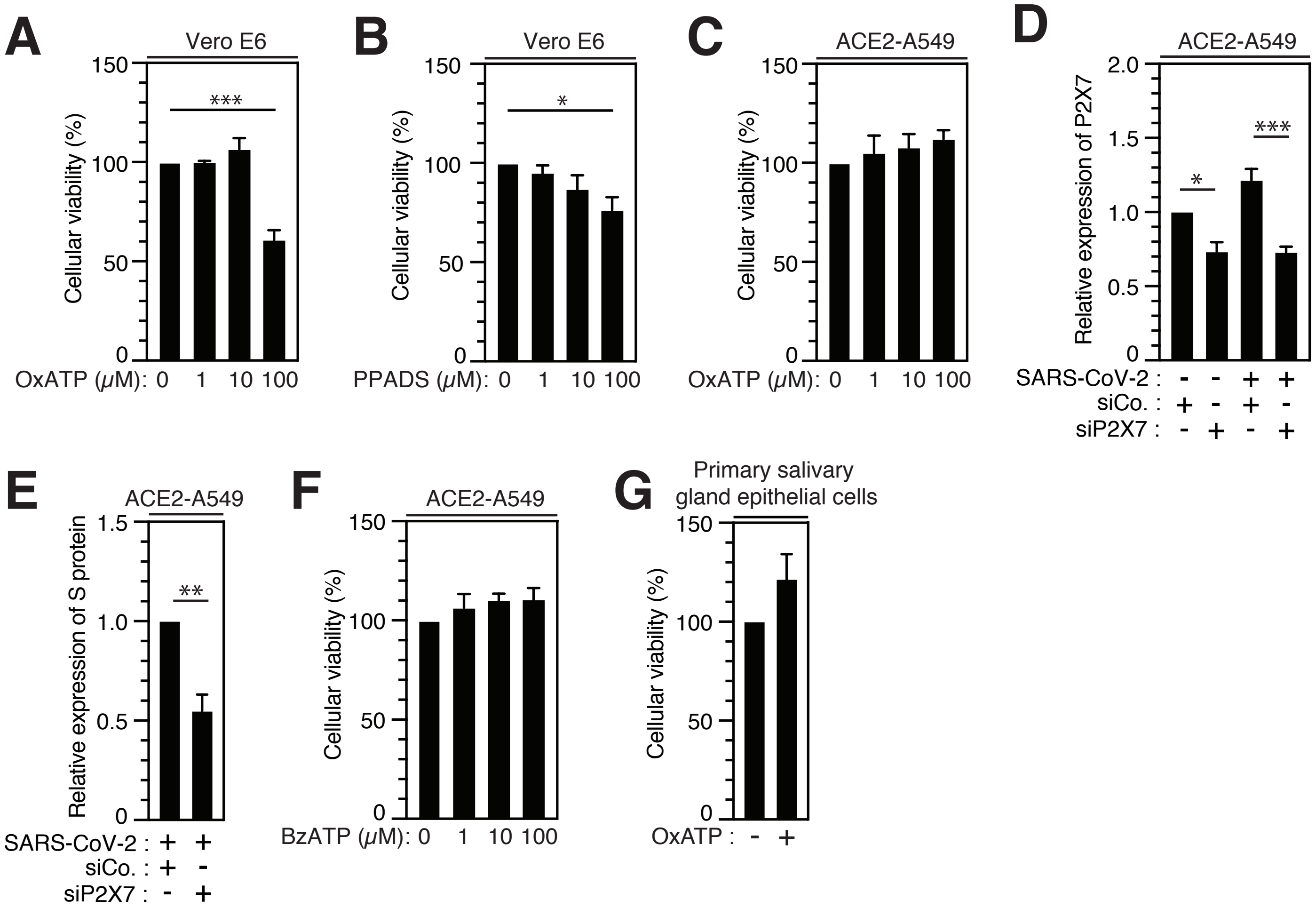
A
Vero E6
Cellular viability (%)
150
100
50
0
***
OxATP (μM): 0 1 10 100
B
Vero E6
Cellular viability (%)
*
PPADS (μM): 0 1 10 100
C
ACE2-A549
Cellular viability (%)
OxATP (μM): 0 1 10 100
D
ACE2-A549
Relative expression of P2X7
2.0
1.5
1.0
0.5
0
*
***
SARS-CoV-2 : - - + +
siCo. : + - + -
siP2X7 : - + - +
E
ACE2-A549
Relative expression of S protein
1.5
1.0
0.5
0
**
SARS-CoV-2 : + +
siCo. : + -
siP2X7 : - +
F
ACE2-A549
Cellular viability (%)
BzATP (μM): 0 1 10 100
G
Primary salivary gland epithelial cells
Cellular viability (%)
OxATP : - +

**LECUYER#S4**

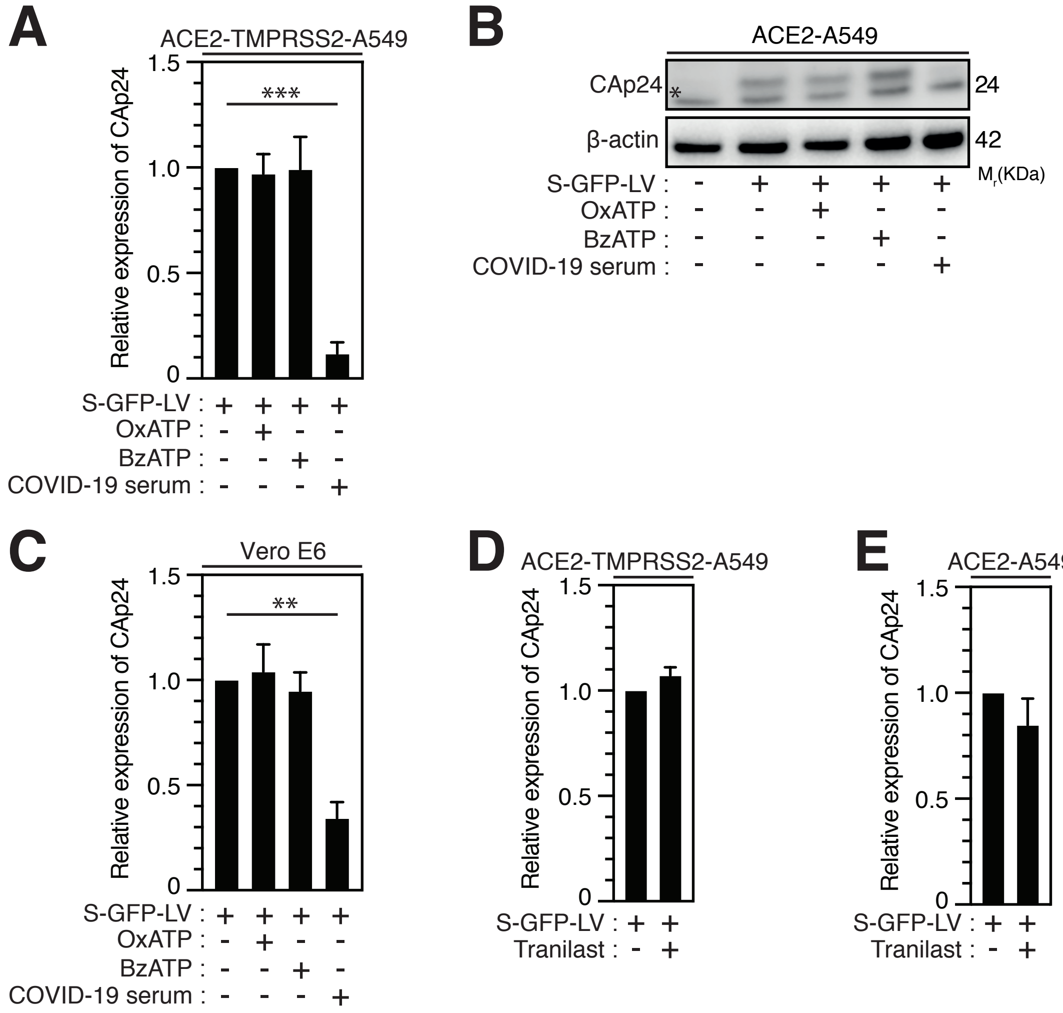
A
ACE2-TMPRSS2-A549
Relative expression of CAp24
1.5
1.0
0.5
0
***
S-GFP-LV : + + + +
OxATP : - + - -
BzATP : - - + -
COVID-19 serum : - - - +
B
ACE2-A549
CAp24
*
24
β-actin
42
Mr(KDa)
S-GFP-LV : - + + + +
OxATP : - - + - -
BzATP : - - - + -
COVID-19 serum : - - - - +
C
Vero E6
Relative expression of CAp24
1.5
1.0
0.5
0
**
S-GFP-LV : + + + +
OxATP : - + - -
BzATP : - - + -
COVID-19 serum : - - - +
D
ACE2-TMPRSS2-A549
Relative expression of CAp24
1.5
1.0
0.5
0
S-GFP-LV : + +
Tranilast : - +
E
ACE2-A549
Relative expression of CAp24
1.5
1.0
0.5
0
S-GFP-LV : + +
Tranilast : - +

**LECUYER#S5**

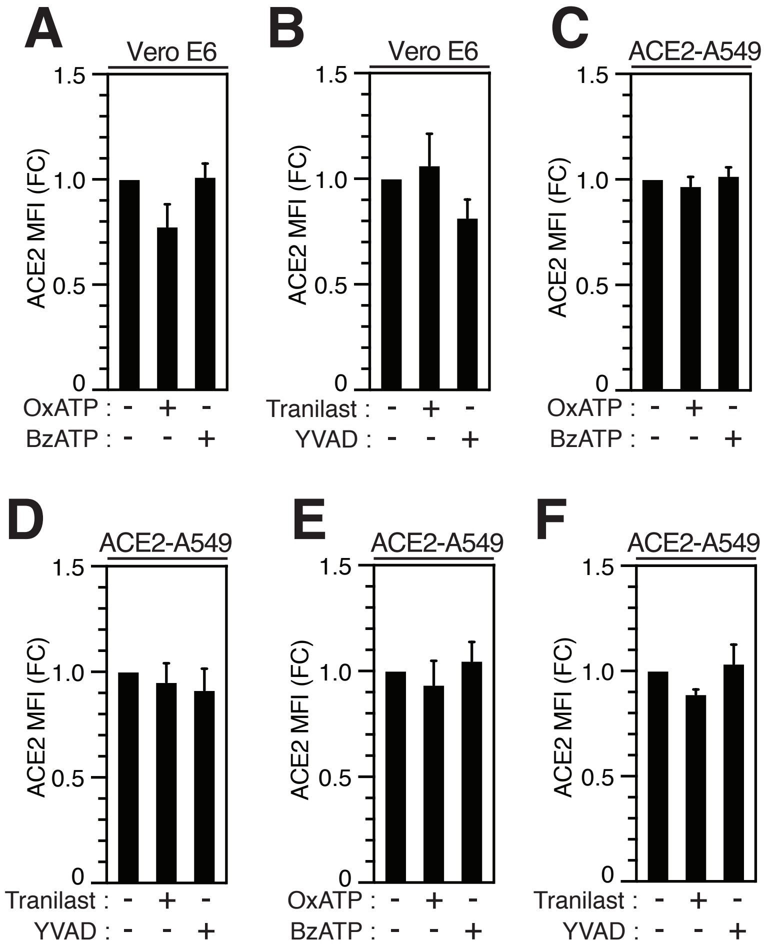
A
Vero E6
ACE2 MFI (FC)
1.5
1.0
0.5
0
OxATP : - + -
BzATP : - - +
B
Vero E6
ACE2 MFI (FC)
1.5
1.0
0.5
0
Tranilast : - + -
YVAD : - - +
C
ACE2-A549
ACE2 MFI (FC)
1.5
1.0
0.5
0
OxATP : - + -
BzATP : - - +
D
ACE2-A549
ACE2 MFI (FC)
1.5
1.0
0.5
0
Tranilast : - + -
YVAD : - - +
E
ACE2-A549
ACE2 MFI (FC)
1.5
1.0
0.5
0
OxATP : - + -
BzATP : - - +
F
ACE2-A549
ACE2 MFI (FC)
1.5
1.0
0.5
0
Tranilast : - + -
YVAD : - - +

**LECUYER#S6**
